# Waterborne, abiotic and other indirectly transmitted (W.A.I.T.) infections are defined by the dynamics of free-living pathogens and environmental reservoirs

**DOI:** 10.1101/525089

**Authors:** Miles D. Miller-Dickson, Victor A. Meszaros, Francis Baffour-Awuah, Salvador Almagro-Moreno, C. Brandon Ogbunugafor

**Affiliations:** Department of Ecology and Evolutionary Biology – Brown University, Providence RI, 02906 USA; Department of Mathematics and Statistics – University of Vermont, Burlington, VT, 05405 USA; Burnett School of Biomedical Sciences – University of Central Florida, Orlando, FL, 32827 USA; National Center for Integrated Coastal Research – University of Central Florida, Orlando, FL, 32816 USA

## Abstract

While the ecology of infectious disease is a rich field with decades worth of empirical evidence and theory, there are aspects that remain relatively under-examined. One example is the importance of the free-living survival stage of certain pathogens, and especially is cases where they are transmitted indirectly between hosts through an environmental reservoir intermediate. In this study, we develop an integrated, broadly applicable mathematical method to examine diseases fitting this description—the waterborne, abiotic and other indirectly transmitted (W.A.I.T.) infection framework. To demonstrate its utility, we construct realistic models of two very different epidemic scenarios: cholera in a densely populated setting with limited access to clean drinking water and hepatitis C virus in an urban setting of injection-drug users. Using these two exemplars, we find that the W.A.I.T. model fortifies the centrality of reservoir dynamics in the “sit and wait” infection strategy, and provides a way to simulate a diverse set of intervention strategies.

## I. INTROODUCTION

Ecology and evolutionary biology have provided a theoretical basis for understanding how interactions between pathogens and their environment shape epidemics. When combined with quantitative modeling methods, it offers a full systems perspective that has helped to characterize the actors, forces and interactions that create infectious diseases [1]–[9]. Classically, the pathogen-host interaction is the presumptive central determinant of infectious disease, and consequently, the focus of modeling efforts: understand it, model it carefully, and one gains a picture for how epidemics arise and persist.

These methods have been successful in balancing simplicity with generality, and have spawned different classes of models, summarized both in terms of the particular mathematical instruments applied (e.g. discrete-time, continuous models, network models, etc) and the particular biologies of host-pathogen systems (e.g. sexually-transmitted, vectorborne disease, food-borne pathogen, etc) [5], [6]. These methods have been effective in many cases, supported by dozens of examples where they have captured the essential character or dynamics of an epidemic [6], [8]. While these existing classifications have served an effective organizational and pedagogical purpose, there remains room for growth in how we translate certain epidemic phenomenon into theoretical formalism.

A concept that has been the object of recent inquiry includes infections transmitted indirectly between hosts via a surface or reservoir intermediate—often abiotic—where the pathogen lives freely and independently of a host [10]–[27], sometimes described as “sit and wait” infections [28]. Much of this research has concentrated on fomite, aerosol and airborne-transmitted viruses [10], [11], [13], [14], [16], [19], [22] and waterborne infections [12], [17], [23], [26], [29]–[33]. Other studies have focused on systems where pathogens are growing in the environment [18], or have explored indirectly-transmitted infections in theoretical terms [21], [24]. One framework for studying indirect, environmental transmission—the environmental infection transmission systems (E.I.T.S)—is engineered with explicit constraints that render its application necessarily narrow [15]. We offer that these prior treatments are individual examples of a general phenomenon that would benefit from a mathematical treatment that is both more rigorous and more broadly applicable. This new formalism should accommodate a wider-range of pathogens than have been previously considered and should emphasize how the environment can comprise multiple discrete dynamic entities (not unlike how host populations are often modeled).

In this study, we develop the “waterborne, abiotic and other indirectly transmitted” (W.A.I.T. or WAIT) paradigm, a generalized framework for understanding “sit and wait” pathogens that are indirectly transmitted through an environmental reservoir intermediate. As the WAIT perspective specifically focuses on the peculiar dynamics of environmental compartments, we argue that it is imbued with more features and applies to a broader set of examples than prior treatments. To demonstrate its novelty and range of application, we fully examine its common relevance to two otherwise disparate modern diseases: cholera and hepatitis C virus. The study proceeds in stages: (1) we first introduce a purely theoretical iteration of WAIT using a standard epidemiological model, explaining how to conceptualize an infection in terms of the WAIT framework, and deriving the reproductive number (*R*_0_) using analytical methods. (2) We apply the WAIT perspective to a waterborne disease that has been the object of several modeling exercises: the transmission of *Vibrio cholerae* in a densely populated setting with limited access to clean drinking water. (3) We then apply the WAIT perspective to a completely different, less-explored disease case: an HCV epidemic among injection drug users (IDU) in a modern urban setting. These constitute different epidemics in terms of pathogen type (*Vibrio cholerae* is a Gram-negative bacterium, HCV a single-stranded RNA virus) and setting. Nonetheless, we conceptualize and model each epidemic scenario, highlighting mathematical features that are unique to the WAIT framework. We discuss how this lens provides new perspective on cholera outbreaks and especially the modern HCV epidemic.

## II. INITIALIZING WAIT: AN ELEMENTARY ADAPTED S-I-R ITERATION

### A. Description

While the emphasis of our examination will reside in how we analyze several modern epidemic systems with the WAIT framework, for explanatory purposes we will begin by describing how it modifies very basic concepts in a classic, purposefully prosaic susceptible-infected-recovered (S-I-R or SIR) mathematical model. While this is analogous to prior methods used to discuss indirect or environmental transmission [17], a full appreciation of both cases discussed in this study (cholera and HCV) would benefit from a full, transparent explanation of the WAIT model-building process. The method is defined by modeling changes in a population of susceptible hosts (“S”), infected hosts (“I”) and recovered (“R”). Classically, flow through the system is defined by contact between susceptible and infected individuals, often driven by a *β* factor, or transmission coefficient.

The WAIT framework would apply to a scenario where the individuals in such a system become infected through an environmental intermediate. Figure 1 is a compartmental model that depicts this interaction, with the two [W] (for “WAIT”) compartments influencing (dashed line) the flow of hosts from the susceptible to infected compartments.

**Fig. 1:**
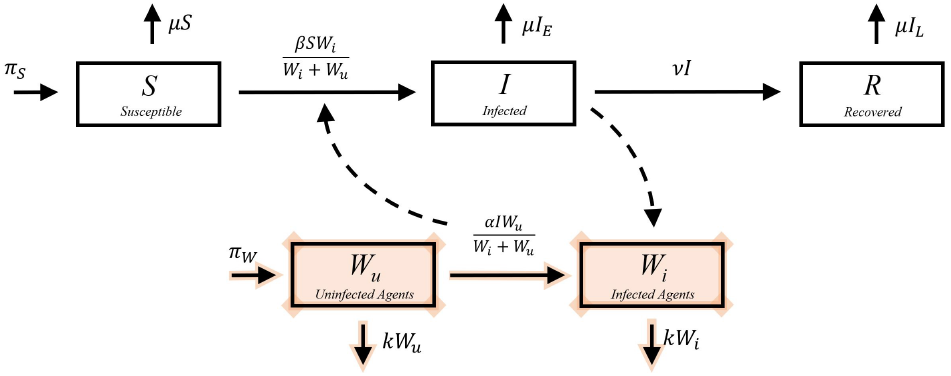
*Adapted SIR* compartmental diagram. This depicts a standard SIR style compartmental model with the added compartments (shaded) corresponding to the WAIT environment. Note the dynamical properties of the *W_i_* and *W_u_* compartments. It is these dynamics that set the WAIT perspective apart from others: environments are often dynamical systems, with an ecology of their own.

### B. The adapted SIR compartmental diagram

The *S*, *I*, and *R* compartments represent the usual *susceptible, infected*, and *recovered* populations of hosts. *W_u_* and *W_i_* represent uninfected and infected populations of *environmental* hosts, respectively.

In traditional SIR models, the rate of new infection (arrow from the *S* compartment to the *I*) is generally proportional to the product of the susceptible and the infected populations, i.e. proportional to *SI*. In the WAIT framework, the environmental compartment, and not a host, drives the rate of infection. In this specific example, the *W_i_* compartment drives the infection such that the rate of infection is proportional to *SW_i_*.

The epidemic is then driven by a series of interactions: interactions between uninfected (susceptible) hosts *S* and the infected (transmitting) environmental compartment *W_i_* and interactions between infected individuals *I* and the uninfected environmental compartment *W_u_*. The epidemic is sustained through infected hosts *I* depositing pathogen into the environmental intermediate, creating new infection in the reservoir, which can then interact with and infect more susceptible hosts *S*, in a process resembling a feedback loop. The dynamics of such a system as we have chosen to model them are captured by the set of dynamical equations below and can be visualized in Figure 1. A derivation of the terms in the model can be read in the Supplemental Appendix.

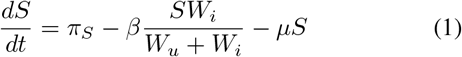

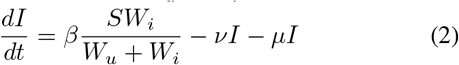

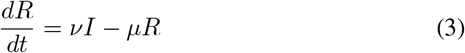

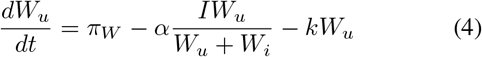

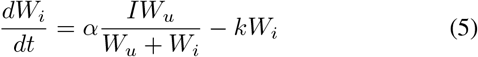

### C. Analytic Equations

Equations 1–5 define the prosaic SIR model. *π_S_* is the birthrate of new susceptible hosts in the environment and *μ* is the fractional death rate of hosts. In this context, *β* represents the *strength* of the interaction between the susceptible hosts *S* and the environment. This will generally be proportional to the rate of contact between the two, and will generally include a factor in it that accounts for the probability that an exposure event actually leads to a new infected host. Similarly, *α* characterizes the strength of interaction between infected hosts *I* and the environmental reservoir; it is also generally proportional to the contact rate between the two and will contain a factor accounting for the probability that when an environmental agent is exposed to infection, it will render the agent infected. *ν* represents the fractional recovery rate, *π_W_* is the birthrate of new uninfected environmental agents, and *k* is the fractional death rate of the environmental agents.

### D. WAIT framework influences the basic reproductive number in a standard SIR model

As an example of how the WAIT model provides a new take on traditional concepts, we briefly consider how the value of the basic reproductive ratio *R*_0_ in this model compares to its SIR counterpart. While *R*_0_ can have different theoretical formulations, we rely on definitions as provided by Jones (2007) [34] and Diekmann et al. 2009 [35]. In a density-dependent SIR model with constant birth of susceptible hosts *π_S_* and death rate proportional to the host population −*μS*, the *R*_0_ value is given by:

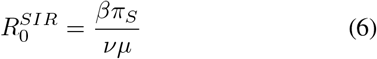

or sometimes, more simply, 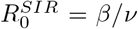, depending on the form of the SIR equations used, e.g. density-dependent, frequency-dependent, constant population, etc. *β* in this equation is the traditional transmission coefficient. In this classic case, it represents the coupling strength between two non-environmental agents, and not one between the environment and the traditional hosts, as in the WAIT iteration. *π_S_*, *μ*, and *ν* have the same interpretation as in our model.

Alternatively, the value of *R*_0_ for the WAIT iteration takes the form:

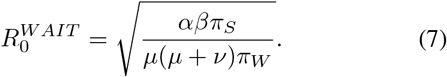

The *R*_0_ formulae (equations 6–7) highlight differences between the models: The square root in the WAIT version arises as a consequence of implementing two infected agents (*I* and *W_i_*) into the model, as opposed to just one in the SIR case. Next, one notices that the *β* factor in the SIR formula is augmented by the additional factor *α* in the WAIT formula, representing a kind of shared dependence between the couplings controlling the *I*-interaction (*α*) and the *S*-interaction (*β*), with the environment. Analogously, what was the responsibility of *π_S_* in the SIR formula is now a shared dependence on *π_S_*/*π_W_*, the ratio of the birthrate of susceptible hosts to that of uninfected environmental agents.

In this case, the two appear as a ratio under the square root, as opposed to a product in the *αβ* case, indicating that these parameters have opposite effects on the value of *R*_0_: when *π_W_* is increased, *R*_0_ decreases, but when *π_S_* is increased, *R*_0_ increases. Whereas, both the *α* and *β* terms will increase the value of *R*_0_ when increased. This presentation allows one to disaggregate the dependencies of the disease burden, as parameterized by *R*_0_, between *hosts* and *environmental agents*.

An additional observation can be made regarding the form of the *R*_0_ in the WAIT iteration. 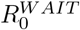 can be viewed as a geometric mean of two other *R*_0_-like values: (1) number of new host infections caused infected environmental agents, and (2) number of new environmental agent infections caused by infected hosts. The first of these *R*_0_-like quantities can be calculated using the rate equations above (1–5), assuming that the system is near the disease free equilibrium (DFE)—the regime where *R*_0_ is calculated. One will find that the rate of new host infections caused by infected environmental agents, near the DFE, is given by *βπ_S_*/(*μπ_W_*). The second *R*_0_-like quantity refers to the number of new environmental agent infections caused by infected hosts, near the DFE. One will find that this value is given by, *α*/(*μ*+*ν*). Thus, one can see that the form of *R*_0_ given in equation 7 can be written as

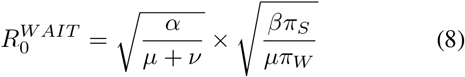

The details of the full calculation can be found in the Supplemental Appendix. Thus, the full *R*_0_ for the WAIT system represents a kind of average of the associated *R*_0_ values for the different modes of transmission. By stressing the role of environmental reservoirs, this modelling perspective has the capacity to dissect properties of dynamics that other models may omit.

### E. Basic rules for building a WAIT model

Constructing models using the WAIT framework requires an understanding of the specific biologies of the agents involved in the system of interest. Consequently, the use of the WAIT framework is as much a conceptual exercise as it is a mathematical one, driven by domain expertise on the problem of interest. The implementation of the framework can be summarized by three key concept-activities:

- Identify the relevant agent-compartments that define the dynamics of the system of interest (e.g. S, I, R, latent infected, etc.) and construct a compartmental model that captures interactions between them.
- If infection in the system is driven by interactions between hosts and an environmental reservoir, one can consider and model dynamics of the latter independently. One should ask how pathogens are deposited in this reservoir, and if/how the pathogen of interest replicates and/or survives in this setting. This is a key point of innovation, as it allows the modeler to think carefully about the biology and ecology of the environmental reservoir.
- Consider how the WAIT reservoir interacts with hosts. How does this interaction occur? What parameters would define how effectively pathogens are transmitted into the reservoir? Note that the nature of this reservoir can be : it can be large or small, liquid or solid, mobile or immobile, and there could also be multiple reservoirs. It is generally abiotic in nature, as the transmitting reservoir isn’t a living organism as in vector-borne diseases (one could consider some exceptions, but these descriptions hold for most cases).

Having proposed a brief outline for how to think about and model a disease system using the WAIT perspective, we will apply it to two different, modern epidemic scenarios: first cholera in a densely populated setting with limited access to clean drinking water, and then hepatitis C virus in a community of injection drug users.

## III. THE CHOLERA WAIT MODEL

### A. Description

Here we present a WAIT-style model of cholera (caused by the gram negative bacterium *Vibrio cholerae*), which has been the object of prior modeling efforts [12], [29]–[31], [33]. This specific iteration is based on a simulation of a 2010 outbreak in Haiti [32]. It is set with a population of 400,000 (the approximate population of the Carrefour section of Port Au Prince, Haiti), features seven ordinary differential equations, seven compartmentalized agents, and twenty parameters. It contains high-infectious and low-infectious water reservoirs, which are the mode of transmission [36]. Recent studies have examined other modes of transmission in cholera, (including transmission within households) [37], but we have chosen to focus on waterborne transmission. Treatment and prevention is undertaken via vaccination, antibiotic administration, and modification of the contaminated water dynamics.

### B. Cholera WAIT model: compartmental diagram

The model can be visualized using a compartmental diagram as seen in Figure 2. Some general features, which reflect findings in prior studies [32], [36], [38], include the following:

- Disease is acquired by individuals as they become infected from drinking water from either the low-infectious or high-infectious reservoirs.
- Infected individuals (symptomatic and asymptomatic) shed bacteria into the high-infectious reservoir.
- Bacteria are removed from the system either via decay in the low-infectious reservoir, or via the death of infected individuals. These reservoirs differ in how much *V. cholerae* exist in a given amount of water, with the high-infectious reservoir much more likely to cause disease.
- Some individuals are vaccinated and individuals return to the susceptible compartment as their vaccine induced immunity wanes.
- Individuals enter and exit the system via equal natural birth and death rates, as well as a death rate due to the disease. Since natural birth and death rates are equal, and additional death can occur from disease, there is a loss of susceptible individuals as the model moves forward through time.

**Fig. 2:**
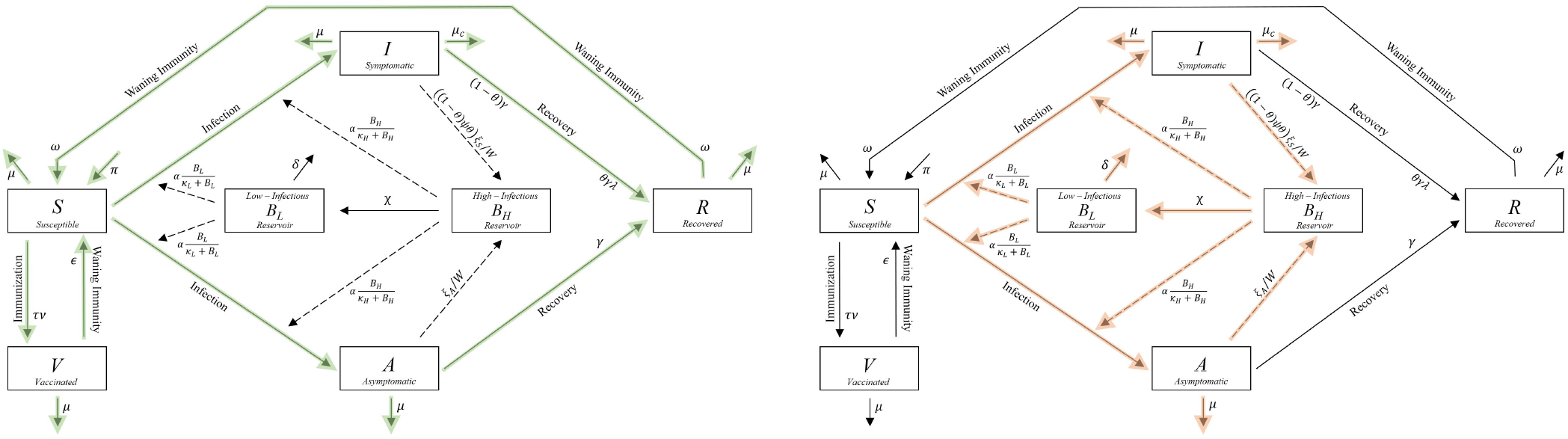
Cholera compartmental diagram. **Left**: Green arrows highlight the flow of the population of hosts through the system. **Right**: here the red arrows highlight flow of disease through the system.

### C. Cholera WAIT model: Analytic equations and parameters

The set of ordinary differential equations (Eq. 9–15) define the dynamics of the system. As outlined in the “elementary adapted S.I.R example,” the environmental dynamics are realized within their own set of differential equations.

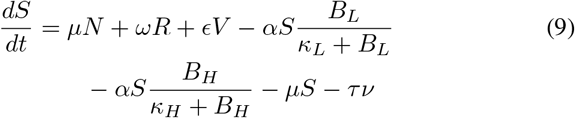

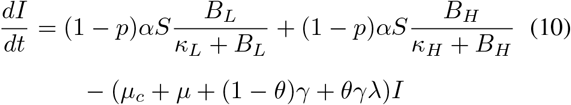

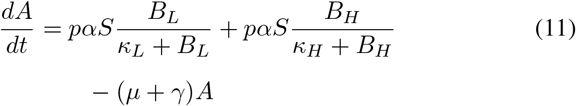

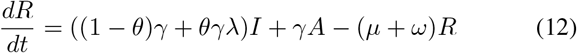

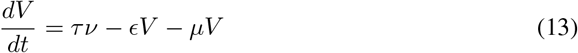

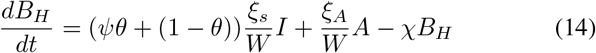

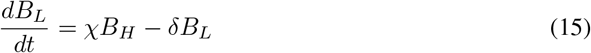

Table I lists the parameters and values used in this simulation. Of note, the 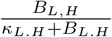 terms in Eq. 9–11 quantify the likelihood that an individual, per day, will be infected with *V. cholerae* given the contaminated water consumption rate *α*. These terms are constructed using the minimum low and high infectious reservoir dose in units of cells per day. Other notable terms include, *α* the water consumption rate and *ξ_s_* the symptomatic individuals excretion rate.

**TABLE I:**
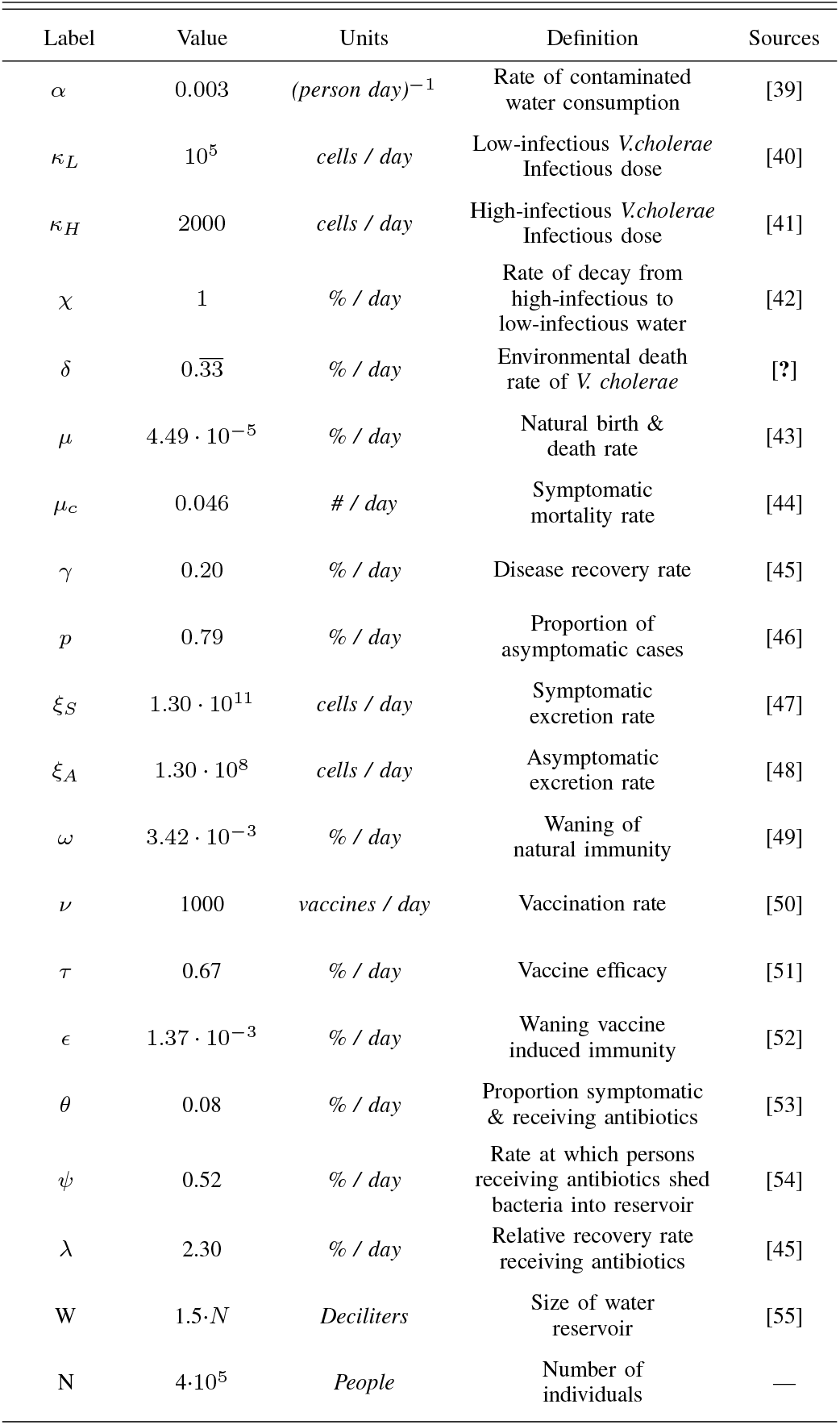
Cholera WAIT model parameters

The initial conditions across all simulations included a population of 400,000 susceptible individuals with an initial small amount of *Vibrio cholerae* present in each reservoir to prime the outbreak (5 cells in each reservoir). Additionally, the values of the symptomatic, asymptomatic, vaccinated, and recovered compartments were set to zero.

### D. Cholera WAIT model: disease dynamics

The simulation was constructed with a time step resolution of 10,000 over a period of 130 days. Figure 3 provides an overview of the dynamics of the disease and the effect of potential interventions. From these simulations, we can observe the behavior of the infected populations (symptomatic and asymptomatic), both of which grow rapidly in the first 20 days of simulation time (Figure 3A). The symptomatic group peaks first, shortly after which the asymptomatic group rises to a higher value. The decay that is seen in the latter days of the outbreak naturally occurs as the total number of susceptible individuals decreases. Further discussion regarding the long term behavior of the model, as well as the susceptible, recovered and vaccinated populations can be found in the Supplemental Appendix.

**Fig. 3:**
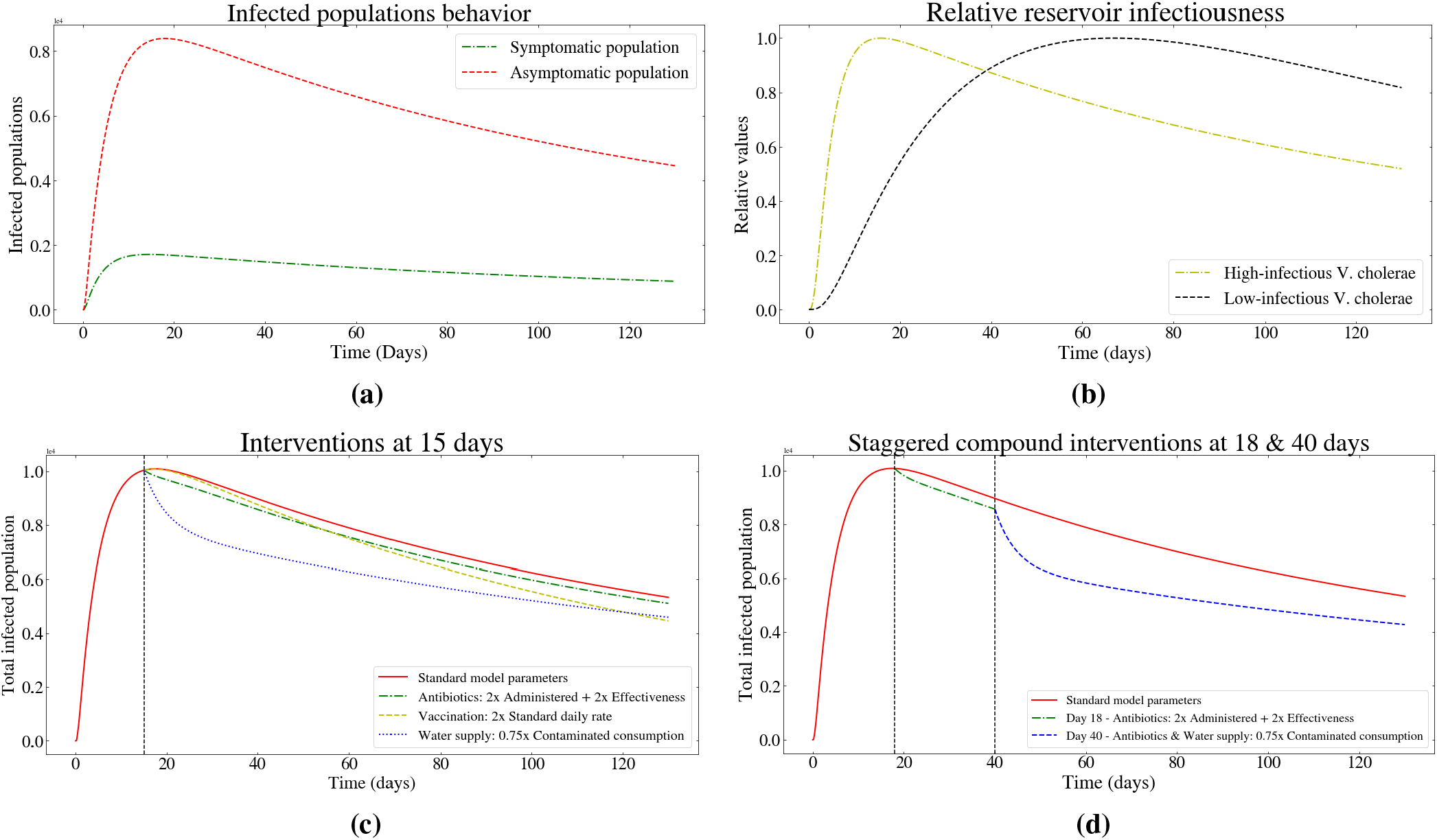
Cholera WAIT model dynamics and simulated interventions. **(a)**: The dynamics of both the symptomatic and asymptomatic infected populations. **(b)**: The fractional relative infectiousness of each reservoir (normalized to 1). **(c)**: Antibiotic, vaccine, and water purification interventions applied at 15 days after the start of the model outbreak **(d)**: Compound staggered interventions mimicking an example real world disease management strategy.

**Fig. 4:**
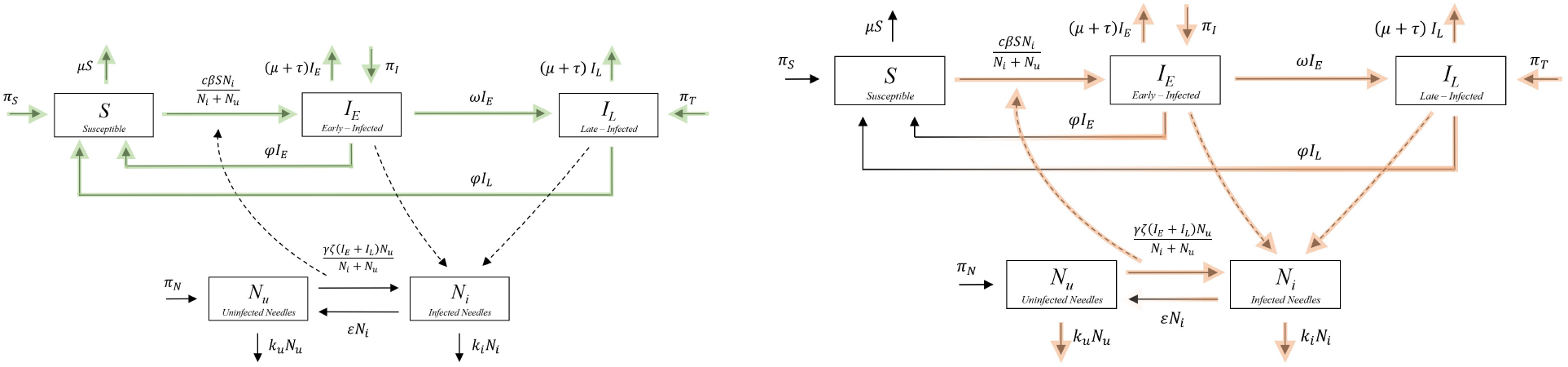
Hepatitis C virus compartmental diagram. **Left:** Green arrows highlight the flow of the population of hosts through the system. **Right:** Red arrows highlight flow of disease through the system, and where there is a color/transparency gradient there is a flow of infection away from an infected compartment towards an uninfected one.

Using the WAIT environmental reservoir equations, one can model the relative changes in water reservoir infectiousness. In Figure 3b, we observe the high-infectious reservoir peak first, driven by the symptomatic population’s shedding of bacteria into the aquatic environmental reservoir. This behavior is is propelled by high excretion rate of symptomatically infected individuals (*ξ_s_*), which is three orders of magnitude larger than (*ξ_A_*). The high-infectious reservoir drives the rise of new infected cases. As the system moves forward in time, the high-infectious reservoir decays via the *χ* parameter, which shifts the disease-driving burden to the low-infectious reservoir.

Vaccination is approximated as a constant number of individuals vaccinated per day, resulting in a linear increase in the vaccinated population. On long time scales (greater than 2500 days) the recovered population peaks and declines as the total number of susceptible individuals ultimately decays, which no longer resembles realistic disease dynamics. Further exploration of this behavior can be found in the Supplemental Appendix.

### E. Cholera WAIT model parameters influence the R_0_

The *R*_0_ provides a signature of the average infectiousness of a given pathogen in a given setting [34], [35]. In this model, given the parameters, **R_0_ = 0.199** (A derivation of the analytic expression for *R*_0_, as well as details regarding the sensitivity analysis, can be found in the Supplementary Appendix.) Thus, in its current configuration, the model does not describe a self-sustaining epidemic. (This is observable through the decay of the asymptomatic and symptomatic cases as depicted in Figure 3a). Additionally, the eigenvalues of the Jacobian matrix for the ODE system at the disease-free equilibrium are real numbers less then or equal to zero, with the largest eigenvalue being zero. Consequently, given enough time, the flow of disease will move to the disease free equilibrium.

The relative sensitivity of *R*_0_ to changes in the model parameters was examined, and we find that *p*, *W*, *α*, and *ξ_S_* are the four most sensitive parameters. These include two parameters strongly related to WAIT aspects of the model: the rate of contaminated water consumption *α*, and the total water reservoir size *W*. Further analysis was also conducted to better understand how adjustments to the model parameters can push *R*_0_ > 1. The results of these analysis can be found in the Supplemental Appendix.

### F. Cholera WAIT model and simulated interventions: antibiotics, vaccination and water purification

Having explained the model structure and analyzed parameter influences on the basic reproductive ratio, we then analyzed a range of potential interventions, comparing their impact on various properties of the cholera dynamics. Specifically, we examine three types of interventions: vaccination, antibiotic administration, and water purification. Each were realized by modifying the respective model parameters that encode information relevant to that intervention.

Figure 3 shows various iterations of the simulation, tracking the sum of both symptomatic and asymptomatic individuals in the standard model (Figure 3a), with various different interventions implemented (3c, 3d). In Figure 3c, we observe how the effect of daily vaccine administration manifests most clearly in the longer term (>100 days) behavior of the dynamics, where it can eventually exceed the effects of reduction in contaminated water consumption (if vaccine effectiveness is sufficiently high). We can also see the large impact of instantaneously reducing contaminated water consumption on the number of infected individuals (in terms of both number of infected individuals, and the rate at which that impact manifests). For example, increasing antibiotic, and vaccine administration two-fold has less immediate impact on disease dynamics than decreasing contaminated water consumption even by a quarter. Together, the cholera WAIT model offers the hypothesis that ideal interventions might include a combination of long-term (e.g. vaccination) and short-term (e.g. water purification) interventions to limit the number of infected cases.

Lastly, compound intervention dynamics are considered (Figure 3d). These aim to simulate disease management strategies where, given practical strategies such as resource allocation and response time, a staggered or combined response is more likely to be implemented. We chose this particular compound intervention to demonstrate how the WAIT model can broadly accommodate the types of modifications that represent public health interventions.

## IV. THE HEPATITIS C VIRUS WAIT MODEL

### A. Description

The HCV model describes a population of approximately 170,000 individuals—based on estimates of the size of the IDU community in New York City [56]—where infected injection drug users may migrate into the population. In this model, needles serve as the environmental intermediate for infection, and the sharing of infected needles will constitute the means of transmitting new infection. Note that while the term “needle” is only one part of a syringe, we use it to refer to the entire syringe. HCV can also be transmitted sexually [57], but in this paper we restrict our attention to transmission through infected needles. As with the cholera model, this section of the main text focuses on the main structure and dynamical properties of the model. Further model details and discussion can be found in the Supplemental Appendix.

### B. HCV WAIT model: Compartmental diagram

We model the dynamics of needle populations, injection drug users, and infected individuals through a series of five ordinary differential equations. The compartments, labeled *S, I_E_, I_L_, N_u_*, and *N_i_* represent the populations of susceptible individuals, early-infected individuals (sometimes referred to as acutely infected), late-stage infected individuals (sometimes referred to as chronically infected), uninfected needles, and infected needles, respectively. This model has several features:

- The susceptible compartment refers to individuals who are injecting drugs and who are sharing needles with other members in the IDU community.
- The needle population is divided into two compartments: infected and uninfected, and we model the dynamics of each compartment separately.
- New infections (of both people and needles) will depend on the fraction of infected or uninfected needles in circulation.
- There are various estimates for the ability of HCV to survive in needles [58] [59]. We incorporate HCV free-living survival via the parameter *ϵ*, which quantifies the rate at which the virus spontaneously clears from infected needles.

### C. HCV WAIT model: Analytic equations and parameters

The dynamics of the HCV transmission process are governed by equations 16–20. The population of individuals who are being treated and those who have recovered are not explicitly modeled in this WAIT iteration, as the dynamics of treatment and recovery are not central to the questions explored in this study. There are, however, several modeling studies of HCV that focus on treatment [60]–[63], and their effects are not ignored in the HCV WAIT model. Entering or leaving treatment (and re-entering the susceptible population, as in case of drug relapse in the IDU population) are approximated by the “birth” terms *π_S_* and *π_T_* and removal terms −*τI_L_* and −*τI_E_*.

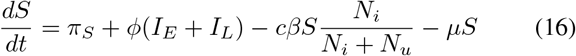

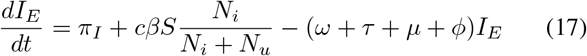

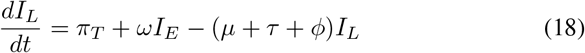

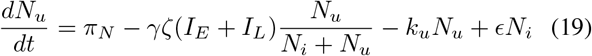

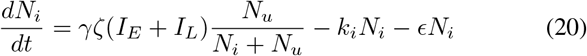

*π_S_* is the birthrate (via migration or first-time use) of new members into a given community of injection drug users (constant in this model). This includes those who may be returning to the population of IDU after recovering from HCV. *π_T_* represents the rate that individuals enter the infected stage coming from an unsuccessful treatment program. *ϕ* represents the daily fraction of individuals infected with HCV who spontaneously clear the infection. *c* represents the rate of sharing a needle with another user (infected or otherwise) per capita. *β* represents the probability that an uninfected individual will convert to HCV+ after injection with an infected needle. *μ* is the combined fractional death and IDU cessation rate (individuals who leave the IDU community). *π_I_* represents the rate of migration of early stage infected individuals into the study population of IDU (assumed to be constant). *ω* is the daily fraction of early-stage infected individuals who progress to the late-stage of infection. *τ* is the daily fraction of infected individuals who go into treatment. *π_N_* is the rate of introduction of uninfected needles into the population of IDU. *γ* is the average number of injections per user per day. *ζ* represents the probability that a new needle becomes infected after use by an infected user. *k_u_* is the daily fraction of uninfected needles which are discarded. *k_i_* is the daily fraction of infected needles which are discarded. Lastly, *ϵ* is the fraction of infected needles which become uninfected in the period of a day due to deactivation (or “death”) of virus populations on the needle. Parameter values and sources can be seen in Table II.

**TABLE II:** HCV WAIT model parameters

| Label | Value | Units | Definition | Sources |
| --- | --- | --- | --- | --- |
| $\pi_S$ | $300 \pm 100$ | person / day | Birthrate of susceptibles (entering the IDU community) | Estimate |
| $\pi_T$ | $25 \pm 10$ | person / day | Rate of treatment failure | Estimate |
| $\phi$ | $0.26 \pm 0.05$ | % / day | HCV self-clearance fraction per day | [60] [67] |
| $c$ | $1 \pm 0.5$ | shares / day | Daily sharing rate of needles per capita | [68] |
| $\beta$ | $0.018 \pm 0.005$ | % | Probability of infection from an infected needle | [69] |
| $\mu$ | $(2.7 \pm 0.5) \cdot 10^{-4}$ | % / day | Fractional rate of removal from IDU community due to cessation & death | [70] |
| $\pi_I$ | $40 \pm 10$ | person / day | Migration rate of new early infected into IDU community | Estimate |
| $\omega$ | $0.5 \pm 0.3$ | % / day | Birthrate of late infections as a fraction of early infections | [71] [72] |
| $\tau$ | $0.04 \pm 0.02$ | % / day | Fractional rate of entering treatments | [73] [74] |
| $\pi_N$ | $(8 \pm 3) \cdot 10^3$ | needles / day | Birthrate of uninfected needles | [75] [76] |
| $\gamma$ | $2.5 \pm 2$ | injections / day | Daily rate of injections per capita | [72] [76] |
| $\zeta$ | $0.25 \pm 0.1$ | % | Probability of transferring infection to a needle | [77] |
| $k_u$ | $0.3 \pm 0.2$ | % / day | Fractional discard rate of uninfected needles | Estimate |
| $k_i$ | $0.3 \pm 0.2$ | % / day | Fractional discard rate of infected needles | Estimate |
| $\epsilon$ | $0.96 \pm 0.03$ | % / day | Fractional decay rate of HCV infection in needles | [59] |

### D. HCV WAIT model parameters influence the R_0_

Having constructed and elaborated on the details of the HCV WAIT model, we now explore how parameters related to the environment (in this case, those framing the population of infected needles) influence the *R*_0_. We directly measured the influence of parameters on the *R*_0_ by considering the effect of modifying the parameters. The value of *R*_0_ was calculated using established methods [34], [35], and is also found to conform to a geometric mean of two other *R*_0_-like quantities, just as in our adapted SIR model in section II. The full calculation and its presentation as a geometric mean is outlined in the Supplemental Appendix.

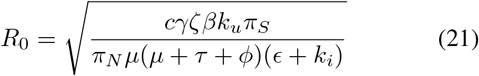

From the form of *R*_0_ given in equation 21, one could surmise that several parameters in the formula have comparable sensitivities. We tested this directly by calculating the Partial Rank Correlation Coefficient (PRCC) for each of the parameters in the model, with respect to *R*_0_, based on methods used in prior studies [64]. We find that parameters related to an interaction with the environmental reservoir (the population of needles) such as *k_u_* (the fractional discard rate of uninfected needles), *γ* (the rate of daily injections per capita), and *π_N_* (the birthrate of new needles) are as central to HCV dynamics as traditionally considered parameters, such as *π_S_* (the birthrate of susceptibles), *c* (the contact or sharing rate of IDU), or *μ* (the combined death and cessation rate of IDU) (Figure 5). This fortifies the notion that WAIT-specific properties dictate the spread of HCV, providing opportunities to explore more precise targeting by public health interventions.

**Fig. 5:**
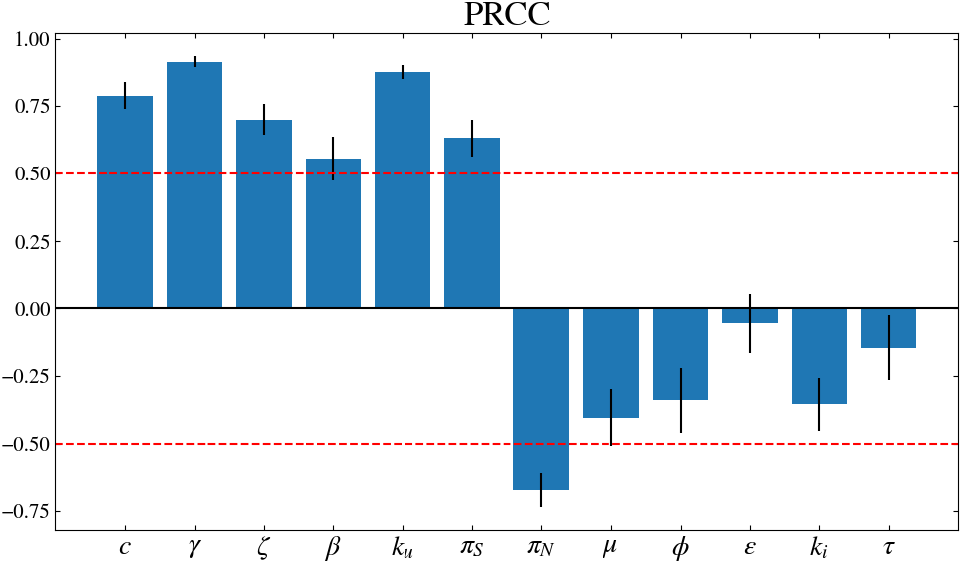
R_0_ sensitivity in HCV: the Partial Rank Correlation Coefficient (PRCC). A PRCC calculation was performed for *R*_0_ using Latin Hypercube Sampling. Parameters were sampled from uniform distributions with widths specified by the ranges given in Table II. The PRCC calculation was repeated for 50 independent iterations. The average of these iterations is shown here, with the standard deviations for each parameter shown as the error bars.

### E. HCV WAIT model and simulated interventions: needle-exchange programs

Having demonstrated the structural relevance of the WAIT framework in terms of how it influences the basic reproductive number, we can consider the utility of the model with respect to other properties, including how it offers insight into potential interventions.

One such intervention may be the implementation of needle-exchange programs. Needle-exchange programs are an example of “harm reduction” public health strategies that aim to reduce harm stemming from behaviors that put the affected individuals or communities at risk [65]. These policies are controversial, but have been demonstrated to be effective interventions for HIV and HCV in certain settings [66]. With respect to the HCV WAIT model, some of these programs (especially ones targeting injection equipment, like safe injection sites) can increase the discard rate of infected needles by providing a safe location to use and discard needles, while also providing uninfected needles to IDU. In our model, parameters like the needle discard rate, *k_i_* and *k_u_*, and *π_N_* are affected by needle exchange programs. Figure 6 demonstrates how *R*_0_ is affected by these parameters. One can see that *R*_0_ can be reduced by increasing *k_i_*—the infected needle discard rate—along a fixed value of *π_N_*—the birthrate of uninfected needles—and that increasing *π_N_* along a fixed value of *k_i_* has the same effect. It is also evident that *R*_0_ can be reduced more rapidly by increasing *k_i_* and *π_N_* simultaneously, as expected. In this way, the proportion of infected needles is reduced because of an increase in clean needles and a reduction of infected ones, lowering *R*_0_.

**Fig. 6:**
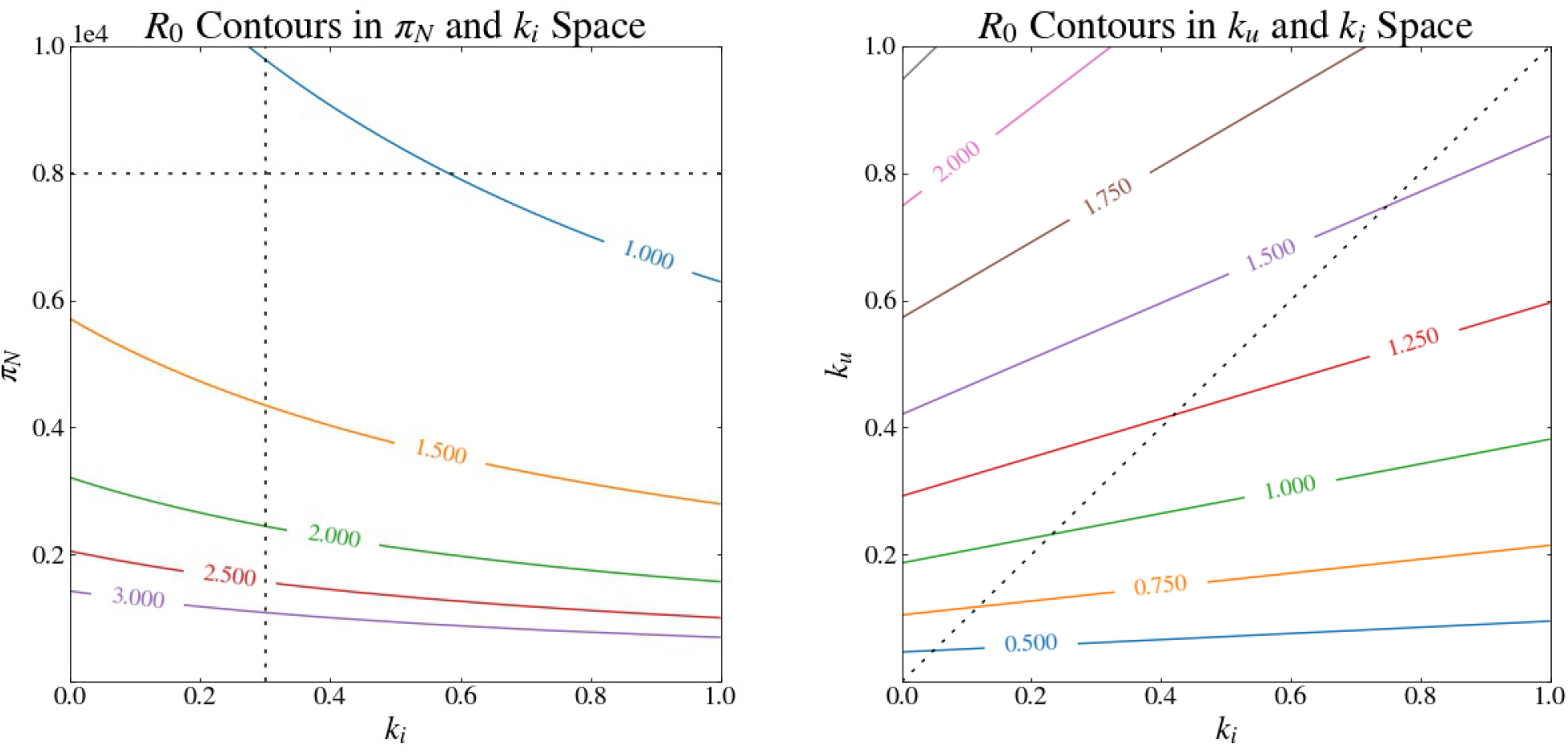
HCV R_0_ as a function of various model features. **Left**: The relationship between the rate of acquisition of clean needles *π_N_* and the discard rate of infected needles *k_i_* with respect to various values of *R*_0_. The curves are contours of constant values of *R*_0_ and are labelled as such. The vertical and horizontal dashed lines indicate the chosen values for their respective parameters. **Right**: The relationship between the infected and uninfected needle discard rate, with respect to *R*_0_. The diagonal line represents when *k_u_* = *k_i_*. Notice that moving upwards along this diagonal increases *R*_0_.

In Figure 6, we demonstrate how changing *k_u_* and *k_i_* modifies the value of *R*_0_. Notice that *R*_0_ is reduced by increasing *k_i_* across fixed values of *k_u_*. The opposite effect is observed when increasing *k_u_* along fixed values of *k_i_*. That is, removing infected needles at an increased rate may decrease infection risk in a population of IDU, while doing the opposite can increase the risk. One can also see that increasing *k_u_* and *k_i_* simultaneously along the dashed line—where *k_u_* = *k_i_*—will increase *R*_0_. This indicates that if a distinction between infected and uninfected needles cannot be established in a given setting, then it may be more helpful to add a higher proportion of uninfected needles than it would be to remove a higher proportion of all needles.

## V. DISCUSSION

In this study, we propose a framework—the waterborne, abiotic and other indirectly transmitted pathogen paradigm (WAIT)—for modeling infectious diseases where transmission is dictated by an interaction between hosts and dynamic multi-compartment environmental reservoirs. We use it to explore two very different modern epidemics of significant public health consequence: cholera and hepatitis C virus. We first establish it through a simple form, an elementary adapted SIR formulation, and highlight how it modifies fundamental properties of epidemics such as the basic reproductive number (*R*_0_). We then build mathematical models to explore cholera and HCV disease dynamics. We demonstrate how the integrated framework highlights potentially overlooked properties of these systems, and highlights potential avenues for intervention. The model is compatible with existing canon in epidemiology, and can accommodate theory in the ecology and evolution of infectious disease. For example, the “curse of the pharaoh hypothesis,” which predicts that pathogens with high free-living survival can be more virulent [78], can be modeled on a population scale using the WAIT framework. In addition, its careful treatment of environmental dynamics makes it amenable to questions regarding how changes in climate, social policy, or behavior may influence the spread of disease.

In the context of cholera, modeling the aquatic reservoir allows us to interrogate the interplay between several notable aspects of cholera epidemics. For example, the interaction between the relative size of the body of water and changes in infected populations, and the notion that the symptomatic and asymptomatic infectious populations fuel different aspects of the epidemic (initial and longer-term phase, respectively). These are areas of present and future inquiry.

In the context of HCV, the WAIT perspective reveals that the dynamics of injection needle populations are crucial to the spread of HCV, which offers a lens on approaches that might attenuate outbreaks of HCV. To the best of our knowledge, no existing mathematical model of HCV epidemiology has interrogated the dynamics in this manner. It is our hope that the HCV modeling community exploits and improves upon this perspective by further adapting the model to study HCV in specific settings.

More generally, The WAIT framework is driven by a contextual understanding of the disease dynamics, where the model parameterization follows the intuition of the scientist. Consequently, it broadens the scope of diseases that can be responsibly understood using mathematical or computational models, potentially generates hypotheses, and fosters improved recapitulation (and possibly prediction) of epidemics driven by “sit and wait” pathogens.

## ACKNOWLEDGMENTS

The authors would like to thank B.Linas for input on the parameters. The authors would like to thank J.Andrews, I.Diakite, S.Robinson, S.Scarpino for various contributions on the topic.

## CODE AVAILABILITY

Code for the mathematical models presented in this manuscript are available on GitHub (https://github.com/OgPlexus).

## Notes

https://github.com/OgPlexus/WAIT

## REFERENCES

[1] Roy M Anderson and Robert M May. Population biology of infectious diseases: Part i. Nature, 280(5721):361, 1979.

[2] Robert M May and Roy M Anderson. Population biology of infectious diseases: Part ii. Nature, 280(5722):455, 1979.

[3] Roy M Anderson and Robert M May. Infectious diseases of humans: dynamics and control. Oxford university press, 1992.

[4] Herbert W Hethcote. The mathematics of infectious diseases. SIAM review, 42(4):599–653, 2000.

[5] Matt J Keeling and Pejman Rohani. Modeling infectious diseases in humans and animals. Princeton University Press, 2011.

[6] Sarah P Otto and Troy Day. A biologist’s guide to mathematical modeling in ecology and evolution. Princeton University Press, 2011.

[7] Sanjay Basu and Jason Andrews. Complexity in mathematical models of public health policies: a guide for consumers of models. PLoS medicine, 10(10):e1001540, 2013.

[8] Eric T Lofgren, M Elizabeth Halloran, Caitlin M Rivers, John M Drake, Travis C Porco, Bryan Lewis, Wan Yang, Alessandro Vespignani, Jeffrey Shaman, Joseph NS Eisenberg, et al. Opinion: Mathematical models: A key tool for outbreak response. Proceedings of the National Academy of Sciences, 111(51):18095–18096, 2014.

[9] Joshua S Weitz. Quantitative viral ecology: dynamics of viruses and their microbial hosts, volume 73. Princeton University Press, 2016.

[10] Elliot C Dick, Lance C Jennings, Kathy A Mink, Catherine D Wartgow, and Stanley L Inborn. Aerosol transmission of rhinovirus colds. Journal of Infectious Diseases, 156(3):442–448, 1987.

[11] F Xavier Abad, Rosa M Pinto, and Albert Bosch. Survival of enteric viruses on environmental fomites. Applied and environmental microbiology, 60(10):3704–3710, 1994.

[12] Cláudia Torres Codeço. Endemic and epidemic dynamics of cholera: the role of the aquatic reservoir. BMC Infectious diseases, 1(1):1, 2001.

[13] Stephanie A Boone and Charles P Gerba. Significance of fomites in the spread of respiratory and enteric viral disease. Applied and Environmental Microbiology, 73(6):1687–1696, 2007.

[14] Thomas P Weber and Nikolaos I Stilianakis. Inactivation of influenza a viruses in the environment and modes of transmission: a critical review. Journal of infection, 57(5):361–373, 2008.

[15] Sheng Li, Joseph NS Eisenberg, Ian H Spicknall, and James S Koopman. Dynamics and control of infections transmitted from person to person through the environment. American journal of epidemiology, 170(2):257–265, 2009.

[16] Raymond Tellier. Aerosol transmission of influenza a virus: a review of new studies. Journal of the Royal Society Interface, 6(suppl_6):S783–S790, 2009.

[17] Joseph H Tien and David JD Earn. Multiple transmission pathways and disease dynamics in a waterborne pathogen model. Bulletin of mathematical biology, 72(6):1506–1533, 2010.

[18] Majid Bani-Yaghoub, Raju Gautam, Zhisheng Shuai, P Van Den Driessche, and Renata Ivanek. Reproduction numbers for infections with free-living pathogens growing in the environment. Journal of biological dynamics, 6(2):923–940, 2012.

[19] Jijun Zhao, Joseph E Eisenberg, Ian H Spicknall, Sheng Li, and James S Koopman. Model analysis of fomite mediated influenza transmission. PloS one, 7(12):e51984, 2012.

[20] Romulus Breban. Role of environmental persistence in pathogen transmission: a mathematical modeling approach. Journal of Mathematical Biology, 66(3):535–546, 2013.

[21] Michael H Cortez and Joshua S Weitz. Distinguishing between indirect and direct modes of transmission using epidemiological time series. The American Naturalist, 181(2):E43–E52, 2013.

[22] N Van Doremalen, T Bushmaker, and VJ Munster. Stability of middle east respiratory syndrome coronavirus (mers-cov) under different environmental conditions. Eurosurveillance, 18(38):20590, 2013.

[23] Meili Li, Junling Ma, and P van den Driessche. Model for disease dynamics of a waterborne pathogen on a random network. Journal of mathematical biology, 71(4):961–977, 2015.

[24] Thomas Caraco, Carrie A Cizauskas, and Nang Wang. Environmentally transmitted parasites: Host-jumping in a heterogeneous environment. Journal of theoretical biology, 397:33–42, 2016.

[25] Andrew F Brouwer, Marisa C Eisenberg, Justin V Remais, Philip A Collender, Rafael Meza, and Joseph NS Eisenberg. Modeling biphasic environmental decay of pathogens and implications for risk analysis. Environmental science & technology, 51(4):2186–2196, 2017.

[26] Andrew F Brouwer, Mark H Weir, Marisa C Eisenberg, Rafael Meza, and Joseph NS Eisenberg. Dose-response relationships for environmentally mediated infectious disease transmission models. PLoS computational biology, 13(4):e1005481, 2017.

[27] Joanne P Webster, Anna Borlase, and James W Rudge. Who acquires infection from whom and how? disentangling multihost and multi-mode transmission dynamics in the ‘elimination’era. Phil. Trans. R. Soc. B, 372(1719):20160091, 2017.

[28] Bruno A Walther and Paul W Ewald. Pathogen survival in the external environment and the evolution of virulence. Biological Reviews, 79(4):849–869, 2004.

[29] E Bertuzzo, R Casagrandi, M Gatto, I Rodriguez-Iturbe, and A Rinaldo. On spatially explicit models of cholera epidemics. Journal of the Royal Society Interface, 7(43):321–333, 2009.

[30] Lorenzo Mari, Enrico Bertuzzo, Lorenzo Righetto, Renato Casagrandi, Marino Gatto, Ignacio Rodriguez-Iturbe, and Andrea Rinaldo. Modelling cholera epidemics: the role of waterways, human mobility and sanitation. Journal of the Royal Society Interface, 9(67):376–388, 2011.

[31] Ashleigh R Tuite, Joseph Tien, Marisa Eisenberg, David JD Earn, Junling Ma, and David N Fisman. Cholera epidemic in haiti, 2010: using a transmission model to explain spatial spread of disease and identify optimal control interventions. Annals of internal medicine, 154(9):593–601, 2011.

[32] Jason R Andrews and Sanjay Basu. Transmission dynamics and control of cholera in haiti: an epidemic model. The Lancet, 377(9773):1248–1255, 2011.

[33] Marisa C Eisenberg, Zhisheng Shuai, Joseph H Tien, and P Van den Driessche. A cholera model in a patchy environment with water and human movement. Mathematical Biosciences, 246(1):105–112, 2013.

[34] James Holland Jones. Notes on r0. California: Department of Anthropological Sciences, 2007.

[35] O. Diekmann, J. A. P. Heesterbeek, and M. G. Roberts. The construction of next-generation matrices for compartmental epidemic models. Journal of the Royal Society Interface, 7:873–885, 2010. doi:10.1098/rsif.2009.0386.

[36] Salvador Almagro-Moreno and Ronald K Taylor. Cholera: environmental reservoirs and impact on disease transmission. Microbiology spectrum, 1(2), 2013.

[37] Christine Marie George, Khaled Hasan, Shirajum Monira, Zillur Rahman, KM Saif-Ur-Rahman, Mahamud-ur Rashid, Fatema Zohura, Tahmina Parvin, Md Sazzadul Islam Bhuyian, Md Toslim Mahmud, et al. A prospective cohort study comparing household contact and water vibrio cholerae isolates in households of cholera patients in rural bangladesh. PLoS neglected tropical diseases, 12(7):e0006641, 2018.

[38] Hartley DM, Morris JG Jr, Smith DL. Hyperinfectivity: A critical element in the ability of v. cholerae to cause epidemics? PLOS Medicine, 3(1):e7, 2005. https://doi.org/10.1371/journal.pmed.0030007.

[39] Kolaye, G., Bowong, S., Houe, R., Aziz-Alaoui, M., & Cadivel, M. Mathematical assessment of the role of environmental factors on the dynamical transmission of cholera. 2018, August 23. https://www.sciencedirect.comhttps://doi.org/10.1016/j.cnsns.2018.06.023.

[40] Ayeni A. O. Domestic Water Source, Sanitation and High Risk of Bacteriological Diseases in the Urban Slum: Case of Cholera in Makoko, Lagos, Nigeria. Journal of Environment Pollution and Human Health, 2(1):12–15, 2005. http://www.sciepub.com/reference/32827.

[41] Ashfaqul Alam, Regina LaRocque, Jason Harris, Cecily Vanderspurt, Edward Ryan, Firdausi Qadri, and Stephen Calderwood. Hyperinfectivity of human-passaged vibrio cholerae can be modeled by growth in the infant mouse. Infection and Immunity, 73(10):6674–6679, 2005. https://iai.asm.org/content/73/10/6674.

[42] Merrell, D. S., Butler, S. M., Qadri, F., Dolganov, N. A., Alam, A., Cohen, M. B., Camilli, A. Host-induced epidemic spread of the cholera bacterium. Nature, 2002, June 06. https://www.nature.com/articles/nature00778.

[43] Department of Economic and Population Division Social Affairs. World population prospects the 2008 revision, highlights, working paper. 2008. http://www.un.org/esa/population/publications/wpp2008/wpp2008highlights.pdf.

[44] Ministère de la santédu cameroun, données choléra. 2014.

[45] Rahaman, M. M., Majid, M. A., Alam, A. K., & Islam, M. R. Effects of doxycycline in actively purging cholera patients: A double-blind clinical trial. Antimicrobial Agents and Chemotherapy, 10(4):610–612, 1976. DOI:10.1128/AAC.10.4.610.

[46] Benenson, A. S., Mosley, W. H., Fahimuddin, M., Oseasohn, R. O. Cholera vaccine field trials in east pakistan. 2. effectiveness in the field. 38(3):359–372, 1968. http://www.sciepub.com/reference/32827.

[47] J. G. & L K., Morris. Cholera. 1995, January 1. https://cmr.asm.org/content/8/1/48.

[48] Levine, M. M., Kaper, J. B., Herrington, D., Losonsky, G., Morris, J. G., Clements, M. L. Hall, R. Volunteer studies of deletion mutants of vibrio cholerae o1 prepared by recombinant techniques. 1988, January. https://www.ncbi.nlm.nih.gov/pmc/articles/PMC259251/.

[49] Woodward, W. E., Mosley, W. H., & McCormack, W. M. The spectrum of cholera in rural east pakistan. i. correlation of bacteriologic and serologic results. 1970, May. https://www.ncbi.nlm.nih.gov/pubmed/4912066.

[50] Amber Hsiao, Sachin N Desai, Vittal Mogasale, Jean-Louis Excler Laura Digilio. Lessons learnt from 12 oral cholera vaccine campaigns in resource-poor settings. Bulletin of the World Health Organization, (95):303–312, 2017.

[51] Thiem, V. D., Deen, J. L., Von, L., Canh, D. G., Anh, D. D., Park, J. K., Clemens, J. D. Longterm effectiveness against cholera of oral killed wholecell vaccine produced in vietnam. 2006, May 15. https://www.ncbi.nlm.nih.gov/pubmed/16580760.

[52] Lopez A. L. Kanungo S. Paisley A. Manna B. Ali M. Clemens J. D Sur, D. Efficacy and safety of a modified killed-whole-cell oral cholera vaccine in india: An interim analysis of a cluster-randomized, doubleblind, placebo-controlled trial. 2009, November 14. https://www.ncbi.nlm.nih.gov/pubmed/1981900.

[53] Hartley DM, Morris JG Jr, Smith DL. Cholera and Post Earthquake Response in Haiti. 1(18):1–9, 2011.

[54] W. A. Khan. Single-dose azithromycin for childhood cholera. 2010, July 23. https://link.springer.com/article/10.1007/s13312-010-0054-x.

[55] Water resources assessment of haiti. 1999, August. https://www.gvsu.edu/cms4/asset/CE59D505-D565-6FFC-FF0D167C69F76EFC/acoewaterresourcesofhaiti.pdf.

[56] Don C Des Jarlais, Theresa Perlis, Samuel R Friedman, Sherry Deren, Timothy Chapman, Jo L Sotheran, Stephanie Tortu, Mark Beardsley, Denise Paone, Lucia V Torian, et al. Declining seroprevalence in a very large hiv epidemic: injecting drug users in new york city, 1991 to 1996. American journal of public health, 88(12):1801–1806, 1998.

[57] Norah A. Terrault, Jennifer L. Dodge, Edward L. Murphy, John E. Tavis, Alexi Kiss, T. R. Levin, Robert G. Gish, Michael P. Busch, Arthur L. Reingold, Miriam J. Alter. Sexual transmission of hepatitis c virus among monogamous heterosexual couples: The hcv partners study. Hepatology, 57(3):881–889, 2013.

[58] Needle Stick Injuries in the Community. 13(3):205–18, 2008.

[59] Paintsil, E., He, H., Peters, C., Lindenbach, B. D., & Heimer, R. Survival of hepatitis c virus in syringes: Implication for transmission among injection drug users. The Journal of Infectious Diseases, 202(7):984–990, 2010. http://doi.org/10.1086/656212.

[60] Vickerman P, Martin N, Turner K, Hickman M. Can needle and syringe programmes and opiate substitution therapy achieve substantial reductions in hepatitis c virus prevalence? model projections for different epidemic settings. Addiction, 107:1984–1995, 2012.

[61] Martin NK, Pitcher AB, Vickerman P, Vassall A, Hickman M. Optimal control of hepatitis c antiviral treatment programme delivery for prevention amongst a population of injecting drug users. PLoS ONE, 6(8), 2011. doi:10.1371/journal.pone.0022309.

[62] Fraser H, Martin NK, Brummer-Korvenkontio H, Carrieri P, Dalgard O, Dillon J, Goldberg D, Hutchinson S, Jauffret-Roustide M, Kåberg M, Matser AA, Maticic M, Midgard H, Mravcik V, Øvrehus A, Prins M, Reimer J, Robaeys G, Schulte B, van Santen DK, Zimmermann R, Vickerman P, Hickman M. Model projections on the impact of hcv treatment in the prevention of hcv transmission among people who inject drugs in europe. Journal of Hepetology, 68(3):402–411, 2018. https://www.ncbi.nlm.nih.gov/pubmed/29080808.

[63] Natasha K Martin, Peter Vickerman, Jason Grebely, Margaret Hellard, Sharon J Hutchinson, Viviane D Lima, Graham R Foster, John F Dillon, David J Goldberg, Gregory J Dore, et al. Hepatitis c virus treatment for prevention among people who inject drugs: modeling treatment scale-up in the age of direct-acting antivirals. Hepatology, 58(5):1598–1609, 2013.

[64] S. M. Blower and H. Dowlatabadi. Sensitivity and uncertainty analysis of complex models of disease transmission: An hiv model, as an example. International Statistical Review, 62(2):229–243, August, 1994.

[65] G Alan Marlatt, Mary E Larimer, and Katie Witkiewitz. Harm reduction: Pragmatic strategies for managing high-risk behaviors. Guilford Press, 2011.

[66] Charlotte Van Den Berg, Colette Smit, Giel Van Brussel, Roel Coutinho, and Maria Prins. Full participation in harm reduction programmes is associated with decreased risk for human immunodeficiency virus and hepatitis c virus: evidence from the amsterdam cohort studies among drug users. Addiction, 102(9):1454–1462, 2007.

[67] Micallef JM, Kaldor JM, Dore GJ. Spontaneous viral clearance following acute hepatitis c infection: a systematic review of longitudinal studies. Journal of Viral Hepatitis, 13(1):34–41, 2006. https://www.ncbi.nlm.nih.gov/pubmed/16364080.

[68] Ilias Gountas, Vana Sypsa, Sarah Blach, Homie Razavi, Angelos Hatzakis. Hcv elimination among people who inject drugs. modelling pre- and post–who elimination era. PLOS ONE, 2018. https://doi.org/10.1371/journal.pone.0202109.

[69] Centers for Disease Control and Prevention (CDC). Recommendations for follow-up of health-care workers after occupational exposure to hepatitis c virus. MMWR Morb Mortal Wkly Rep, 46(26):603–606, 1997. https://www.ncbi.nlm.nih.gov/pubmed/9221329.

[70] Hickman M, Hope V, Brady T, Madden P, Jones S, Honor S, Holloway G, Ncube F, Parry J. Hepatitis c virus (hcv) prevalence, and injecting risk behaviour in multiple sites in england in 2004. Journal of Viral Hepatitis, 14(9):645–652, 2007. https://www.ncbi.nlm.nih.gov/pubmed/17697017.

[71] Poynard T, Bedossa P, Opolon P. Natural history of liver fibrosis progression in patients with chronic hepatitis c. The Lancet, 349(9055):825–32, 1997.

[72] Adnan Khan, Sultan Sial, and Mudassar Imran. Transmission Dynamics of Hepatitis C with Control Strategies. The Journal of Computational Medicine, 2014(654050):18, 2014. http://dx.doi.org/10.1155/2014/654050.

[73] Grebely, Jason Conway, B Raffa, J Lai, C Krajden, Mel Tyndall, Mark. 578 uptake of hepatitis c virus (hcv) treatment among injection drug users (idus) in vancouver, canada. Journal of Hepatology, 44(6):214–215, 2006. https://doi.org/10.1371/journal.pone.0202109.

[74] Seal K, Kral A, Lorvick J, Gee L, Tsui J, et al. Among injection drug users, interest is high, but access low to hcv antiviral therapy. Journal of General Internal Medicine, 20:171, 2005.

[75] Gold K. Analysis: the impact of needle, syringe, and lancet disposal on the community. Journal of Viral Hepatitis, 5(4):848–850, 2011. https://www.ncbi.nlm.nih.gov/pubmed/21880224.

[76] Daliah I Heller, Denise Paone, Anne Siegler, Adam Karpati. The syringe gap: An assessment of sterile syringe need and acquisition among syringe exchange program participants in new york city. Harm Reduction Journal, 6(1), 2009. https://www.ncbi.nlm.nih.gov/pmc/articles/PMC2631523/.

[77] Short LJ, Bell DM. Risk of occupational infection with blood borne pathogens in operating and delivery room settings. Am J Infect Control, 21:343–50, 1993.

[78] Masashi Kamo and Michael Boots. The curse of the pharaoh in space: free-living infectious stages and the evolution of virulence in spatially explicit populations. Journal of theoretical biology, 231(3):435–441, 2004.

