## Supplemental Appendix for "Waterborne, abiotic and other indirectly transmitted (W.A.I.T.) infections are defined by the dynamics of free-living pathogens and environmental reservoirs"

Below we provide additional information on several aspects of the project, including discussion points, and various details on the respective models.

#### WAIT Background: Connection to concepts in evolutionary biology and ecology

In addition to epidemiological modeling efforts, there have been robust evolutionary examinations of the causes and consequences of free-living infections. With regards to the evolution of virulence [1], [2], it has been studied in terms of the "Curse of the Pharoah" hypothesis, where a free-living survival stage would directly influence pathogen virulence [3], [4]. In non-disease ecology, the concept of propagule pressure bears resemblance to free-living survival, though the former is discussed in a much different context (e.g. seed dispersal) [5]. All of these perspectives—epidemiology, modeling, evolutionary biology, ecology—demonstrate that scientists from across fields have reflected on how the indirect transmission strategy might have evolved, and how it manifests in modern epidemics.

#### Initializing WAIT: Introductory model

##### *Derivation and justification for terms in the model*

We begin by presenting the equations governing the dynamics of the introductory model, what we refer to in the main text as the elementary adapted SIR model.

$$\frac{dS}{dt} = \pi_S - \beta\frac{SW_i}{W_u + W_i} - \mu S \tag{1}$$

$$\frac{dI}{dt} = \beta\frac{SW_i}{W_u + W_i} - \nu I - \mu I \tag{2}$$

$$\frac{dR}{dt} = \nu I - \mu R \tag{3}$$

$$\frac{dW_u}{dt} = \pi_W - \alpha\frac{IW_u}{W_u + W_i} - kW_u \tag{4}$$

$$\frac{dW_i}{dt} = \alpha\frac{IW_u}{W_u + W_i} - kW_i \tag{5}$$

The framework of this introductory model, and of the WAIT scheme in general, is such that it passes the spread of infection to a tertiary intermediate—the environmental reservoir—before the infection spreads to susceptible hosts. We achieve this paradigm in the model's equations by including a drainage term (-$\beta\frac{SW_i}{W_u+W_i}$) in the $S$ equation which feeds into the $I$ equation. This term replaces one which, in the usual SIR model, is normally proportional to $SI$, i.e. one representing a coupling between the uninfected hosts $S$ and infected hosts $I$. In the introductory WAIT version, the analogous term represents a coupling between uninfected hosts $S$ and infected *environmental agents* $W_i$. In this way, the spread of infection can be seen to depend on the extent of the infection in the reservoir, and on the size the uninfected population of hosts as well. A notable distinction between SIR and WAIT is that the WAIT coupling term is not proportional to $SW_i$ alone, as one might have expected, but rather to $SW_i/(W_u + W_i)$—i.e. to the *fraction* of infected environmental agents in the system. The basis for this decision relies on an assumption that the interaction rate between the environmental reservoir and hosts is *frequency-dependent*, w.r.t. encounters with the reservoir, as opposed to *density-dependent*, in the sense of the total size of the reservoir $W_u + W_i$. This can be clarified by giving a short derivation of the term in question: First, assuming that $\beta$ represents the number of contact incidents between susceptible hosts $S$ and the environmental reservoir per time step per host, then the quantity $\beta S$ is simply the contact rate between susceptible hosts and the environmental reservoir (here we assume that the contact rate with the reservoir is density-dependent w.r.t. the host population size but not w.r.t. the reservoir size $W_u + W_i$). Lastly, the fraction of these contact incidents where a susceptible host comes in contact with the *infected* portion of the environment should be $W_i/(W_u+W_i)$. Thus, the rate of such incidents should be $\beta S \times W_i/(W_u+W_i)$. Here, we assume that it is these contact incidents, i.e. those between susceptible hosts and the infected portion of the environment that give rise to new infection among the host population.

The infection within the environmental reservoir is itself a dynamical system and is governed by the $W_u$ and $W_i$ equations. One will notice that a term similar to the one described above quantifies the rate at which infection spreads in the reservoir. This term ($\alpha\frac{IW_u}{W_u+W_i}$) has behind it analogous logic: Here, $\alpha$ represents the number of contact incidents with the reservoir by *infected* hosts per time step per host. Thus, $\alpha I$ is the number of times an infected host contacts the reservoir per time step. The fraction of these encounters where the infected hosts come in contact with the uninfected portion of the reservoir is $W_u/(W_i + W_u)$, and so the total rate of encounters between an infected host and uninfected environmental agent is $\alpha I \times W_u/(W_i + W_u)$. Other terms in the model are either simple Malthusian-type rates or constant rates.

#### $R_0^{WAIT}$ *as a geometric mean*

We also take the opportunity to describe the value of $R_0$, which was claimed to be the geometric mean of two other $R_0$-like quantities. $R_0$ can be calculated using traditional methods, using, for instance, the *Next-generation Matrix*. Here, we assume that we have done so and have found the form found in equation 7 in the main text. Then, we will show that two of the factors appearing under the square root in the formula can be regarded as $R_0$-like quantities. Namely, there is the number of new *host*-infections caused by infected environmental agents, and there is the number of new *environmental agent*-infections caused by infected hosts—both of which are calculated at the disease free equilibrium (DFE).

The first of these $R_0$-like quantities represents the number of new host infections caused by an infected environmental agent, when the system is near the DFE. We begin by noticing that the rate of new host infections (according to equations 1–5) is given by $\beta SW_i/(W_u + W_i)$. Near the DFE, $S \approx \pi_S/\mu$ and $W_i/(W_u + W_i) \approx W_i/W_u$, where the latter fact results from a Taylor series about the DFE, using that $W_i << W_u$. Near the DFE, $W_u \approx \pi_W/k$ as well. Thus, the rate of new host infections near the DFE, *per infected environmental agent* $W_i$, is given by $\beta\pi_S k/(\mu\pi_W)$ (notice that we divide out by $W_i$ because we are interested in the per capita rate). The *number* of new infections caused by an environmental agent is then this rate times the average amount of time that an environmental agent spends in the infected state. The average amount of time an agent spends in a compartment is given by the reciprocal of the *exit-rate* of the compartment, in this case by $1/k$. Thus, in the product of this rate and $1/k$, the $k$ factor cancels out, and so the number of new host infections in the time that an environmental agent is infected is given by,

$$R_0^A = \frac{\beta\pi_S}{\mu\pi_W}$$

(near the DFE). We use the superscript $A$ to distinguish this $R_0$-like quantity from the other. The second $R_0$-like quantity represents the number of new environmental agent infections, per infected host, that occur in the average amount of time that a host is infected, near the DFE. This number is determined from the rate of new environmental agent infections, $\alpha IW_u/(W_u + W_i)$, in equations 1–5. Near the DFE, the fraction $W_u/(W_u + W_i) \approx 1$, so the rate of new infections of environmental agents, *per infected host*, is given simply by $\alpha$. The average amount of time that a host spends in the infected compartment is given by the reciprocal of the exit-rate, which is given by $1/(\mu + \nu)$. Thus, the number of new infections of environmental agents that occur in the time that a host is infected is given by,

$$R_0^B = \frac{\alpha}{\mu + \nu}$$

Thus, one finds that the $R_0$ equation presented in equation 7 in the main text can be written as,

$$R_0^{WAIT} = \sqrt{\frac{\beta \pi_S}{\mu \pi_W}} \times \sqrt{\frac{\alpha}{\mu + \nu}} = \sqrt{R_0^A R_0^B}$$

### The Cholera WAIT Model

#### *Complete model dynamics & longer timescale behavior*

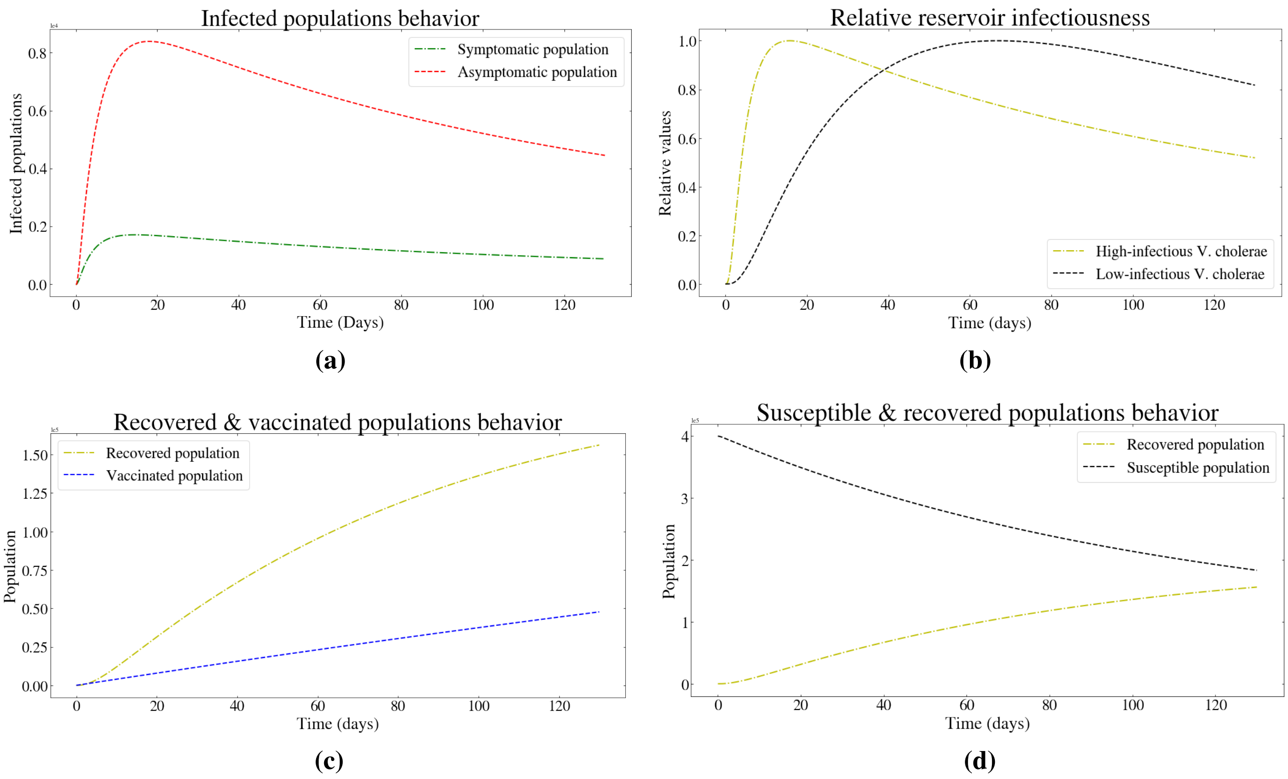


**Fig. S1:** Cholera WAIT model dynamics over 100+ days of integrated time. Of note: panel **(c)** shows the time-linear relationship of the vaccination rate

In order to explore the complete dynamics of the this model over 100+ days of time, simple graphs were produced showing the behavior of every compartment, displayed in the their entirety in Fig. S1.

To elucidate the long time scale behavior of the system, we integrated to a total of 2500 days (approximately 6.85 years) at a resolution of $10^6$ time steps. Here we can see a visual approximation of the system as it approaches equilibrium. We see the effects of the loss of a large amount of the total population as well as the significant decay of *V. cholerae* from the system via the $\delta$ parameter.

Past the $\sim$130 days of simulation time, the model begins to lose real-world relevance. As mentioned in the main text, though this model provides information with regards to the environmental factors that drive a *V. cholera* outbreak, the simulation is self-contained and only models a single idealized cholera outbreak from the beginning to waning. As discussed in the main text, these limitations are commonly observed in differential-equation based epidemiological models.

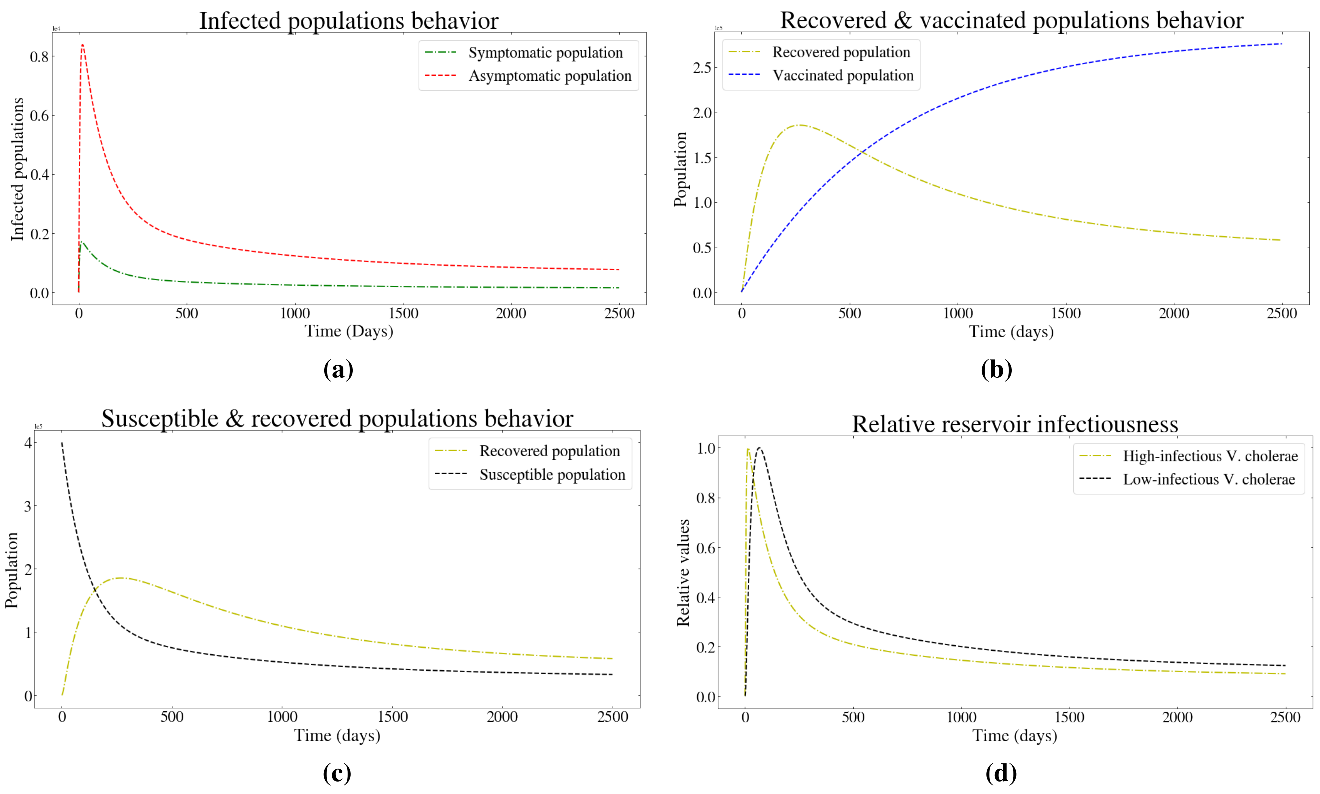


**Fig. S2:** The Cholera model complete long term model dynamics over 2500 days of integrated time, here we can see the effects of the global reduction of susceptible individuals as the model moves past its useful time window

#### *Analytic Derivation of* $\mathrm{R}_0$

The analytic expression for the basic reproductive ratio $R_0$ of *V. cholera* referenced in the main text, and used as part of our sensitivity analysis, was derived following the methods of Diekmann et. al. 2009 [6].

Step (1) of this method is to construct the arrays $f = (f_0, f_1, ..., f_m)$ and $v = (v_0, v_1, ..., v_m)$. The $i^{th}$ element of $f$, denoted $f_i$, is defined to be the rate term associated with the flow of *new* infection *into* the $i^{th}$ compartment. That is, the flow of infection between two infected compartments is not included in $f$. The $i^{th}$ element of $v$, denoted $v_i$, is defined to be the negative of the rate terms associated with all other flows into or out of the $i^{th}$ compartment. That is, the total rate of change of the $i^{th}$ compartment is given by $f_i - v_i$. It turns out that for calculating $R_0$ it suffices to restrict $f$ and $v$ to infected compartments only. In the case of *V. cholerae*, these are the compartments $I$, $A$, $B_L$, and $B_H$.

$$f = \begin{pmatrix} \frac{\alpha(1-p)SB_H}{B_H+\kappa_H} + \frac{\alpha(1-p)SB_L}{B_L+\kappa_L} \\ \frac{\alpha p SB_H}{B_H+\kappa_H} + \frac{\alpha p SB_L}{B_L+\kappa_L} \\ 0 \\ \frac{A\xi_A}{W} + \frac{I(\theta\varphi-\theta+1)\xi_S}{W} \end{pmatrix} \qquad \mathrm{v} = \begin{pmatrix} I\left(\gamma\theta\lambda + \gamma(1-\theta) + \mu_c + \mu\right) \\ A(\gamma+\mu) \\ \delta B_L - B_H\chi \\ \chi B_H \end{pmatrix}$$

Step (2) is to calculate the corresponding $F$ and $V$ matrices. These are $m \times m$ matrices (in this case $m = 4$). They are defined by,

$$F_{ij} = \frac{\partial f_i}{\partial X_j}(x_0)$$
$$V_{ij} = \frac{\partial v_i}{\partial X_j}(x_0)$$

where $X_j$ is the $j^{th}$ agent, selected from the $m$ agents associated with $f$ and $v$, and $x_0$ is the disease-free equilibrium of the model. Calculating these derivatives for the *V. cholera* model one finds,

$$V = \begin{pmatrix} \gamma(1-\theta)+\gamma\theta\lambda+\mu+\mu_c & 0 & 0 & 0 \\ 0 & \gamma+\mu & 0 & 0 \\ 0 & 0 & \delta & -\chi \\ 0 & 0 & 0 & \chi \end{pmatrix}$$

$$F = \begin{pmatrix} 0 & 0 & \frac{(1-p)S\alpha}{B_L+\kappa_L} - \frac{(1-p)S\alpha B_L}{(B_L+\kappa_L)^2} & \frac{(1-p)S\alpha}{B_H+\kappa_H} - \frac{(1-p)S\alpha B_H}{(B_H+\kappa_H)^2} \\ 0 & 0 & \frac{pS\alpha}{B_L+\kappa_L} - \frac{pS\alpha B_L}{(B_L+\kappa_L)^2} & \frac{pS\alpha}{B_H+\kappa_H} - \frac{pS\alpha B_H}{(B_H+\kappa_H)^2} \\ 0 & 0 & 0 & 0 \\ \frac{(\varphi\theta-\theta+1)\xi_S}{W} & \frac{\xi_A}{W} & 0 & 0 \end{pmatrix}$$

the above matrices are still not set to the disease-free equilibria, this is done in the following steps. Step (3) is to calculate the inverse of the matrix $V$. For our case this turns out to be,

$$V^{-1} = \begin{pmatrix} \frac{\gamma\delta\chi+\delta\mu\chi}{\delta(\gamma+\mu)\chi(\gamma(1-\theta)+\gamma\theta\lambda+\mu+\mu_c)} & 0 & 0 & 0 \\ 0 & \frac{\gamma\delta\chi-\gamma\delta\theta\chi+\gamma\delta\theta\lambda\chi+\delta\mu\chi+\delta\mu_c\chi}{\delta(\gamma+\mu)\chi(\gamma(1-\theta)+\gamma\theta\lambda+\mu+\mu_c)} & 0 & 0 \\ 0 & 0 & \frac{1}{\delta} & \frac{1}{\delta} \\ 0 & 0 & 0 & \frac{1}{\chi} \end{pmatrix}$$

Step (4) is to calculate the matrix product $F \cdot V^{-1}$.

$$F \cdot V^{-1} = \begin{pmatrix} 0 & 0 & -\frac{(p-1)S\alpha\kappa_L}{\delta(B_L+\kappa_L)^2} & -\frac{(p-1)S\alpha\left(\delta\kappa_H B_L^2+2\delta\kappa_H\kappa_L B_L+\kappa_L\left(\chi(B_H+\kappa_H)^2+\delta\kappa_H\kappa_L\right)\right)}{\delta\chi(B_H+\kappa_H)^2(B_L+\kappa_L)^2} \\ 0 & 0 & \frac{pS\alpha\kappa_L}{\delta(B_L+\kappa_L)^2} & \frac{pS\alpha\left(\delta\kappa_H B_L^2+2\delta\kappa_H\kappa_L B_L+\kappa_L\left(\chi(B_H+\kappa_H)^2+\delta\kappa_H\kappa_L\right)\right)}{\delta\chi(B_H+\kappa_H)^2(B_L+\kappa_L)^2} \\ 0 & 0 & 0 & 0 \\ \frac{\theta(\varphi-1)\xi_S+\xi_S}{W(\theta(\lambda-1)\gamma+\gamma+\mu+\mu_C)} & \frac{\xi_A}{W\gamma+W\mu} & 0 & 0 \end{pmatrix}$$

Step (5) is to then take $F \cdot V^{-1}$, at the disease-free equilibrium and produce $(F \cdot V^{-1})_{DFE}$,

$$(F \cdot V^{-1})_{DFE} = \begin{pmatrix} 0 & 0 & -\frac{(p-1)\alpha}{\delta\kappa_L} & -\frac{(p-1)\alpha\left(\delta\kappa_H+2\delta\kappa_L\kappa_H+\kappa_L\left(\chi\kappa_H^2+\delta\kappa_L\kappa_H\right)\right)}{\delta\chi\kappa_H^2\kappa_L^2} \\ 0 & 0 & \frac{p\alpha}{\delta\kappa_L} & \frac{p\alpha\left(\delta\kappa_H+2\delta\kappa_L\kappa_H+\kappa_L\left(\chi\kappa_H^2+\delta\kappa_L\kappa_H\right)\right)}{\delta\chi\kappa_H^2\kappa_L^2} \\ 0 & 0 & 0 & 0 \\ \frac{\theta(\varphi-1)\xi_S+\xi_S}{W(\theta(\lambda-1)\gamma+\gamma+\mu+\mu_c)} & \frac{\xi_A}{W\gamma+W\mu} & 0 & 0 \end{pmatrix}$$

Step (6): Lastly, in order to determine the basic reproductive ratio the eigenvalues of $(F \cdot V^{-1})_{DFE}$ are computed, here the largest eigenvalue is given as $R_0$.

$$R_0 = \sqrt{p\left(\xi_A\mu_c + \gamma\left(\theta(\lambda-1)\xi_A + \xi_A + (\theta(1-\varphi)-1)\xi_S\right) + \mu\left(\xi_A + (\theta(1-\varphi)-1)\xi_S\right)\right) + (\gamma+\mu)(\theta(\varphi-1)+1)\xi_S}$$

$$\times \frac{\sqrt{\alpha}\sqrt{\chi\kappa_H\kappa_L + \delta\left(\kappa_L+1\right)^2}}{\sqrt{\delta\chi W(\gamma+\mu)\kappa_H\kappa_L^2\left(\gamma\theta(\lambda-1)+\mu_c+\gamma+\mu\right)}}$$

#### $R_0$ *sensitivity & multi-parameter adjustment*

A sensitivity analysis was conducted in order to determine which parameters have the largest relative effect on the reproductive ratio of the cholera model. Such a ranked analysis of parameter sensitivity allows one to see the added insight gained by introducing a WAIT component to traditional epidemiological models and resolving the environmental dynamics of disease. The stability and robustness of the model can be seen in the long-term behavior of the system and the eigenvalue analysis at the disease-free equilibrium. In the cholera case, there is an added level of complexity present in finding distributions of model parameters that vary monotonically for the Latin Hyper-Cube Sampling algorithm used in generating the PRCC analysis. This is due to the dense nature of this model's analytic expression for $R_0$. For that reason, a partial rank correlation coefficient (PRCC) analysis was forgone.

Of additional relevance: although vaccination is an important part of disease management, no vaccination dynamics (or parameters) appear in the analytic expression for $R_0$ nor the tornado diagram (Figure S3). This is because vaccination is preemptive, and though helpful, is not a characteristic of the basic, essential reproductive capacity of *V. cholerae*.

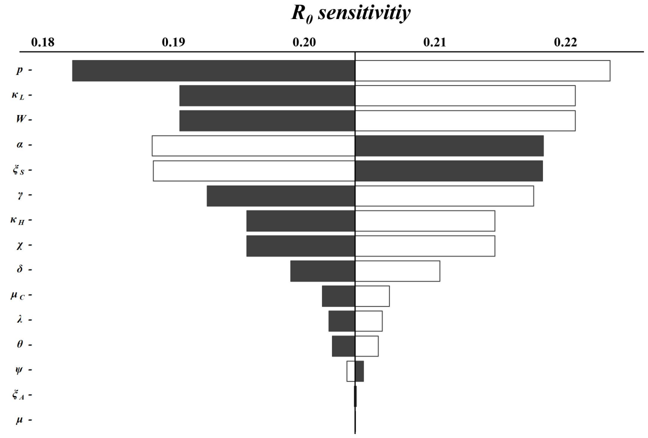


**Fig. S3:** A tornado plot showing the relative sensitivity of $R_0$ to changes in the model parameters black bars indicate the value of $R_0$ when the associated parameter (excluding p) is *increased* by 15% from the value chosen in the model. White bars indicate the value of $R_0$ when the associated parameter (excluding p) is *decreased* by 15%. The proportion of asymptomatic cases, $p$, is hypersensitive in $R_0$, thus in order to better render the diagram $p$ is only symmetrically varied by $\pm 5.5\%$.

In Figure S3, we observe that parameters $W$ and $\alpha$ are the $2^{nd}$ and $3^{rd}$ most sensitive to the reproductive capacity of *V. Cholerae*. Both of these parameters appeal to properties of the aquatic environment and can be influenced by intervention. Within the scope of the model, reducing contaminated water consumption or increasing the size of the aquatic environment (thus diluting the infectious material) reduces its reproductive potential substantially. Furthermore, the effects of these environmental parameters on the reproductive potential of *V. Cholerae* outweighs the impact of non-environmental interventions (antibiotics, vaccines; parameterized in this model by $\theta$, $\lambda$, and $\phi$).

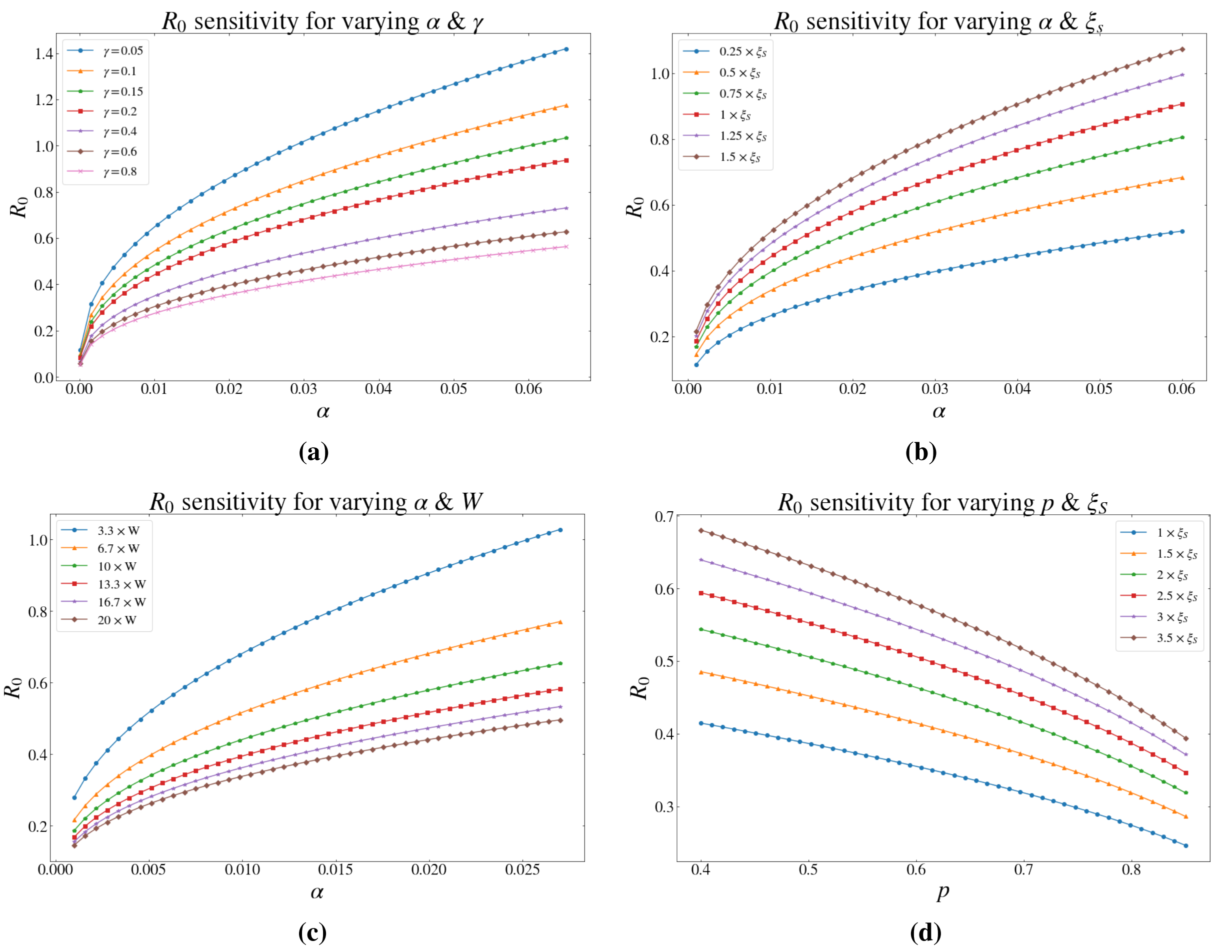


**Fig. S4:** The Cholera model behavior around the space where $R_0 = 1$, as different pairs of parameters are adjusted we can see the outbreak transition into a self-sustaining epidemic

Further analysis was conducted to better understand the regions around certain parameters where this model predicts an epidemic. In Figure S4, we present the results of simultaneous modification of a number of model parameters: $\alpha$ and $\gamma$, $\alpha$ and $\xi_S$, $\alpha$ and $W$, $p$ and $\xi_S$ (the dash lines represent the parameters used in the model).

We can further pontificate on these results in light of several real world hypothetical situations. For example, Figure S4 highlights how $R_0$ increases rapidly as $\gamma$ decreases. In the context of the effects of health and wellness on the body's ability to fight disease, this model suggests that forces that negatively effect overall health would have an effect on disease dynamics. If the recovery rate among infected individuals drops too low (as during a drought or famine) even standard levels of contaminated water consumption may cause an epidemic.

#### *Additional intervention dynamics in Cholera*

Figure S5 shows some additional intervention dynamics that were simulated for the system. Here an additional assortment of starting points and parameter combinations are considered. The interventions shown below were applied in the same manner as discussed in the main text, modifying the driving parameters that define each intervention relative to the nominal model parameters.

As with all the interventions simulated in this model, the staggered intervention starting points are chosen by adjusting the initial conditions to match the conditions of the model at the point of intervention.

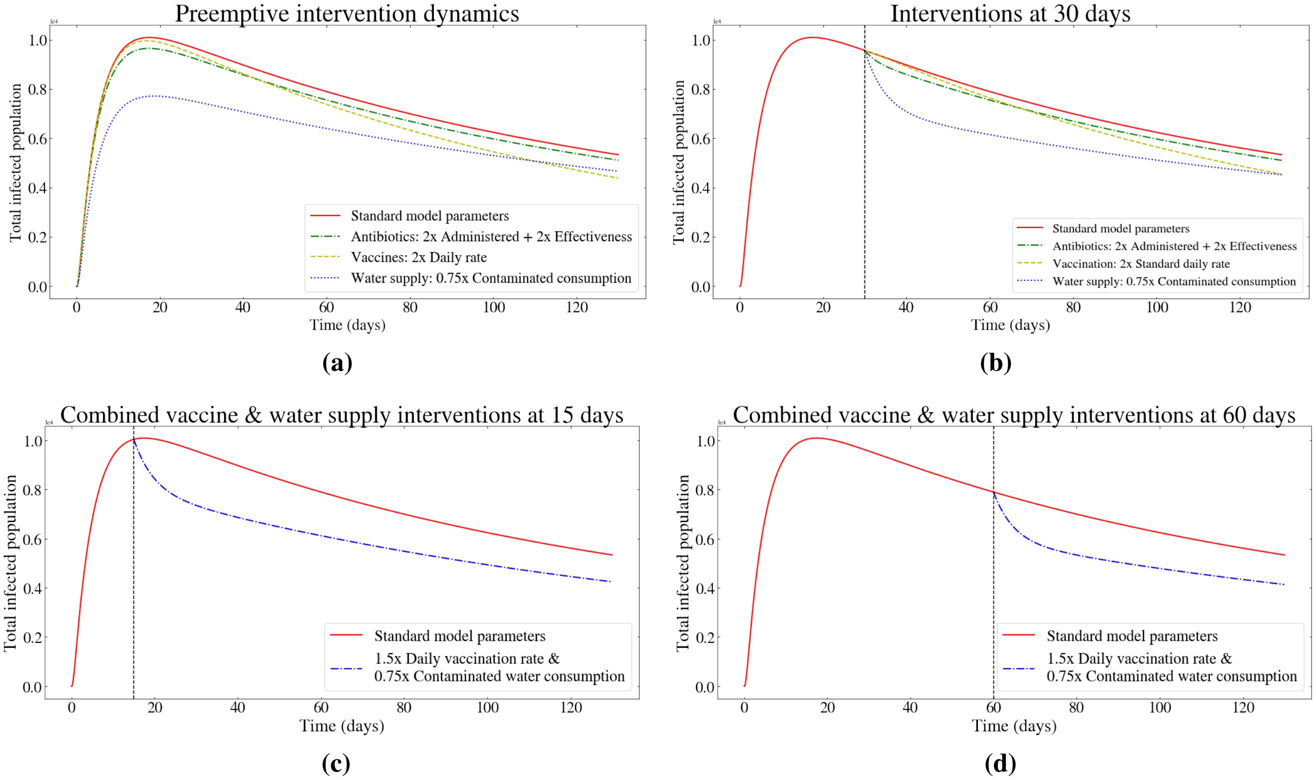


**Fig. S5:** Additional cholera model intervention dynamics highlighting both compound and and individual simultaneous approaches

For preemptive interventions, vaccination or water-treatment can begin before the cholera outbreak begins. This could apply in settings adjacent to ones that have experienced cholera, but have not themselves shown any signs of an outbreak.

Specifically: these intervention dynamics underscore, within the scope of this model, the power of water treatment and the benefit of early interventions early in undermining epidemic dynamics.

#### *Jacobian ODE system analysis & related values*

$$\left(\begin{array}{ccccccc} -\frac{IW}{I+\delta} - \frac{W\theta}{\epsilon+\theta} - \lambda & 0 & 0 & \gamma & V & -\frac{W\delta\tau}{(I+\delta)^2} & -\frac{W\epsilon\tau}{(\epsilon+\theta)^2} \\ -\frac{W}{(I+\delta)(\epsilon+\theta)}(B_L - 1)\left(I(\epsilon+\theta) + \theta(I+\delta)\right) & -\kappa_L\mu_C R + R(\mu_C - 1) - \lambda - \omega & 0 & 0 & 0 & -\frac{W\delta\tau(B_L-1)}{(I+\delta)^2} & -\frac{W\epsilon\tau(B_L-1)}{(\epsilon+\theta)^2} \\ \frac{B_L W(I(\epsilon+\theta)+\theta(I+\delta))}{(I+\delta)(\epsilon+\theta)} & 0 & -R-\lambda & 0 & 0 & \frac{B_L W\delta\tau}{(I+\delta)^2} & \frac{B_L W\epsilon\tau}{(\epsilon+\theta)^2} \\ 0 & R(\kappa_L\mu_C - \mu_C + 1) & R & -\gamma-\lambda & 0 & 0 & 0 \\ 0 & 0 & 0 & 0 & -V-\lambda & 0 & 0 \\ 0 & \frac{S}{B_H}(\chi\mu_C - \mu_C + 1) & \frac{\kappa_H}{B_H} & 0 & 0 & -p & 0 \\ 0 & 0 & 0 & 0 & 0 & p & -A \end{array}\right)$$

Here we present both a symbolic and numeric form of the Jacobian Matrix of the WAIT *V. cholerea* ODE system, the numeric form being at the disease-free equilibrium. The eigenvalues for the disease-free Jacobian are calculated and are all shown to be negative, aside from 0.

$$\begin{pmatrix} -4.49\cdot 10^{-5} & 0.0 & 0.0 & 0.00342 & 0.00137 & -1.5\cdot 10^{-6} & 0.0 \\ 0.0 & -0.2669 & 0.0 & 0.0 & 0.0 & 3.0\cdot 10^{-7} & 0.0 \\ 0.0 & 0.0 & -0.200 & 0.0 & 0.0 & 1.2\cdot 10^{-6} & 0.0 \\ 0.0 & 0.221 & 0.2 & -0.00347 & 0.0 & 0.0 & 0.0 \\ 0.0 & 0.0 & 0.0 & 0.0 & -0.00141 & 0.0 & 0.0 \\ 0.0 & 20834.667 & 21.667 & 0.0 & 0.0 & -1.0 & 0.0 \\ 0.0 & 0.0 & 0.0 & 0.0 & 0.0 & 1.0 & -0.0334 \end{pmatrix}$$

Above we present the numeric form of the DFE Jacobian Matrix, this was calculated using the nominal model parameters applied through the simulation. Below are the eigenvalues corresponding to this Matrix.

$$-0.0035, \quad -0.0014, \quad 0, \quad -0.259, \quad -1.0087, \quad -0.0327, \quad -0.2$$

As with any set of ordinary differential equations the Jacobian Matrix represents the flow of the system in phase space. The eigenvectors give the direction of flow and the sign of the vectors (eigenvalues) orient's them toward or away from an equilibria. Given the Jacobian Matrix is computed at the disease-free equilibria and all its eigenvalues are negative (aside from 0) we can say that the system (given the nominal chosen model parameters) tends to equilibrate and that there is a stable equilibrium at the disease-free equilibrium point in phase space. This is to say that, given the chosen model parameters and an infinite amount of time, the model will move toward and equilibrate at the disease-free equilibrium.

### THE HEPATITIS C VIRUS WAIT MODEL

#### *Additional notes on the structure of the model*

Here we provide justification for two terms in the model that would benefit from elaboration: (a) the rate of new infection of susceptible individuals, $c\beta S\frac{N_i}{N_i+N_u}$, and (b) the rate of new infection of clean needles, $\gamma\zeta(I_E+I_L)\frac{N_u}{N_i+N_u}$; other terms in the model are fairly generic.

For (a), the total incidence rate of receiving needles from other users, per capita, is given by $c$. Then, $cS$ represents the total rate of susceptible hosts receiving needles from other users. In our model, we assume that the likelihood of sharing an infected needle is proportional to the fraction of infected needles in circulation $\frac{N_i}{N_i+N_u}$. Thus, the rate at which susceptible hosts use infected needles is $cS\frac{N_i}{N_i+N_u}$. $\beta$ corresponds to the proportion of such incidences where an injection with an infected needle by an uninfected user leaves the individual infected, and thus the product of the two $c\beta S\frac{N_i}{N_i+N_u}$ represents the rate of new infection among hosts via infected needles.

For (b) (the rate of new infection of clean needles via infected hosts) we suppose that the rate at which needles are infected is proportional to the number of infected individuals and is scaled by the daily rate of injections per capita, $\gamma$. The product, $\gamma(I_E+I_L)$ therefore represents the average number of injections per day by all infected individuals. The proportion $\zeta$ represents the fraction of which that leave a needle infected.

Lastly, to account for the events where an injection drug user can select an already infected needle for an injection event, we multiply by the fraction of needles with the potential for becoming infected, i.e. the uninfected proportion, $\frac{N_u}{N_i+N_u}$. This would most likely manifest as the proportion of uninfected needles in the total population of IDU. In the HCV WAIT model, we approximate this as the proportion of uninfected needles that a particular IDU has access to. This approximation will break down when there is large variance in the distribution of infected and uninfected needles in the infected population of IDU. Note that for the purposes of infecting a needle, we only need to consider the population of infected individuals and thus, do not need to suppose that the distribution of infected and uninfected needles has low variance in the *total* population of IDU.

#### *The dynamics of needles*

Our model allows us to distinguish between the discard rates of infected and uninfected needles. However, while it may be that interventions such as needle-exchange programs are capable of increasing the infected needle discard rate above that of the uninfected needles, in many circumstances, the distinction between the two might be more difficult to disentangle. Thus, in many cases there may not be a discernible distinction between $k_u$ and $k_i$, in which case these rates are effectively equal—we will denote this *universal* discard rate by $k$ $(= k_u = k_i)$.

Our model presents a somewhat counter-intuitive dynamic when $k_u = k_i$: Since the infection rate of needles relies on the proportion of uninfected needles in a population, reducing the proportion of clean needles in the population can actually exacerbate an infection. Namely, increasing the discard rate of all needles will, in particular, reduce the number of uninfected needles in the population but—due to asymmetries in how infected and uninfected needles are treated in the model—can actually lead to an increase in the *proportion* of infected needles in the population, increasing the probability that a user will select an infected needle. We examine the mathematical details of this behavior below.

A public health interpretation of this would simply be removing all needles from the IDU community (through law enforcement, for example) without a proportional increase in safe, clean needles (as supplied in clean-injection sites or needle-exchange programs), potentially intensifying the local HCV epidemic as a result. Figure S6(C) shows the steady state fraction of infected needles in a population of IDU, as a function of this universal discard rate $k$. Notice that for the parameters chosen in our model, the proportion of infected needles increases when the discard rate $k$ is increased.

In such circumstances, the disease burden and value of $R_0$ are both increased. In Figure 5 in the main text, we demonstrate how changing $k_u$ and $k_i$ can modify the value of $R_0$. We find that increasing $k_u$ while moving along a constant value of $k_i$ has the effect of increasing the $R_0$ value. Whereas, increasing $k_i$ while moving along a constant value of $k_u$ reduces $R_0$. That is, removing more uninfected needles while keeping the same infected needle discard rate can harm a population of IDU, and doing the opposite helps the population. One can also see that increasing $k_u$ and $k_i$ simultaneously along the diagonal line—where $k_u = k_i$—will increase $R_0$. This suggests that if a distinction between infected and uninfected needles cannot be determined (as is often the case), then adding clean needles to a population has a larger impact on disease burden than flatly removing all needles.

#### *Equilibrium values of the infected needles*

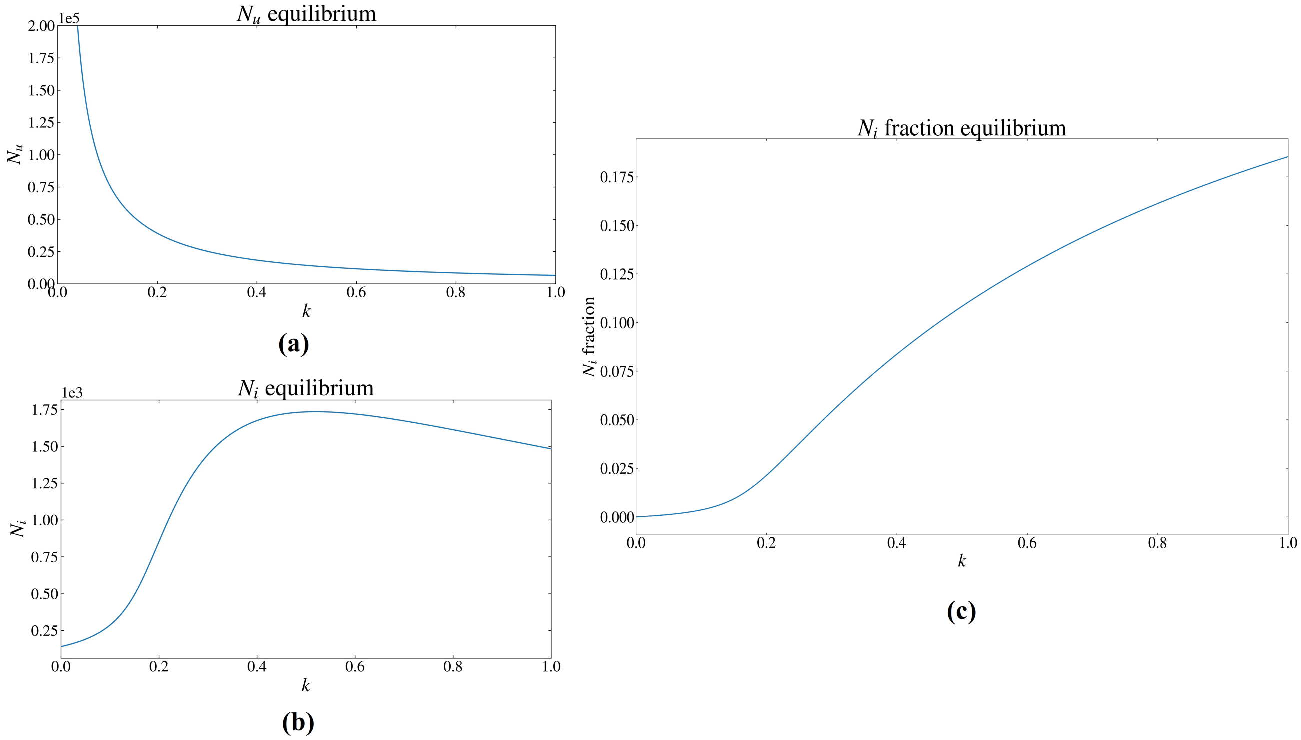


**Fig. S6:** **(a)** represents the number of uninfected needles, $N_u$, as a function of the discard rate, $k$. **(b)** shows the same but for $N_i$, the number of infected needles in the population. **(c)** shows the *fraction* of infected needles in the population across the fractional discard rate $k$, i.e. it shows the value of $N_i/(N_u + N_i)$ at equilibrium as a funciton of $k$. Note, that these are endemic equilibria, whereas the disease-free equilibrium will occur when the infection fully clears from the population of IDUs, and the population of infected needles accordingly goes to zero.

By setting the model equations to zero, one can determine the equilibrium or steady state values of the agents as a function of the parameters in the model. In doing so, we can demonstrate more quantitatively how the proportion of infected and uninfected needles is affected by modifying the discard rate of needles. In Fig. S6 (a) and (b) we plot the equilibrium values for $N_u$ and $N_i$ as a function of $k$ (where $k = k_u = k_i$).

One will notice that the equilibrium value of $N_u$ is a monotonically decreasing function of $k$ (at least for $k$ between $0$ and $1$), whereas $N_i$ actually increases with $k$ first before gradually decreasing. That is, whereas the steady state value of uninfected needles always decreases when the universal discard rate $k$ is increased, the steady state value of the infected needles actually has the potential to rise if $k$ is increased from a sufficiently small value. This is consistent with a prior point about the effects of flatly discarding needles, without regard to their infected or uninfected status: it can increase the proportion of infected needles in a population and thus fuel the epidemic. Fig. S6(C) shows the *fraction* of infected needles in the population, and although one finds that the absolute number of infected needles in equilibrium will subside beyond a certain value of $k$ (in our case this is around $k = 0.5$), we find that the fraction of infected needles is a monotonically increasing function of $k$. That is, for the parameters chosen, discarding needles universally will increase the likelihood of encountering an infected needle in the population of IDUs. We emphasize that this insight was only possible because the needle reservoir (the environment) was modelled separately, a central feature of WAIT.

Below, we provide the results of our analytic calculation of the equilibrium values of *infected needles* as a function of $k$. We find that the infected needle equilibrium can be expressed in the form,

$$N_i^*(k) = \frac{\tilde{a} + \tilde{b}k + \sqrt{\tilde{c} + \tilde{d}k + \tilde{e}k^2}}{\tilde{g}k(\tilde{h}k + \tilde{f})}$$

where the parameters $\tilde{a}, \tilde{b}, \tilde{c}, \tilde{d}, \tilde{e}, \tilde{f}, \tilde{g}$, and $\tilde{h}$ (given explicitly below) are independent of $k$ and which are functions of the other parameters in the model.

From the equations governing the dynamics of infected and uninfected needles, it is evident that the total number of needles $N_u + N_i$ evolves according to the equation:

$$\frac{d(N_u + N_i)}{dt} = \pi_N - k(N_u + N_i)$$

This is determined by adding the equations for $N_u$ and $N_i$ together. Setting this equation to zero shows that the total number of needles in the population, $N_u + N_i$, has as its equilibrium value $\pi_N/k$. From this and the equation provided above for the equilibrium value of infected needles $N_i^*$, one can show that $N_u^*$, the equilibrium value of the uninfected needles, is given by,

$$N_u^*(k) = \frac{1}{k}\left(\pi_N - \frac{\tilde{a} + \tilde{b}k + \sqrt{\tilde{c} + \tilde{d}k + \tilde{e}k^2}}{\tilde{g}(\tilde{h}k + \tilde{f})}\right)$$

Below, we give the explicit form of the parameters in the above formulae:

$$\tilde{a} = -\epsilon\mu\pi_N^2(\mu + \tau + \phi)$$

$$\tilde{b} = \pi_N\left(\gamma\zeta(\beta c(\pi_I + \pi_S + \pi_T) - \mu(\pi_I + \pi_T)) - \mu\pi_N(\mu + \tau + \phi)\right)$$

$$\tilde{c} = \mu^2\pi_N^4\epsilon^2(\mu + \tau + \phi)^2$$

$$\tilde{d} = 2\mu\pi_N^3\epsilon(2c\beta\gamma\zeta(\pi_I + \pi_T)(\mu + \tau) +$$
$$(\mu + \tau + \phi)(-\beta c\gamma\zeta(\pi_I + \pi_S + \pi_T) + \mu(\gamma\zeta(\pi_I + \pi_T) + \pi_N(\tau + \phi)) + \mu^2\pi_N) + 2\beta c\gamma\zeta(\mu + \tau)(\pi_I + \pi_T)))$$

$$\tilde{e} = \pi_N^2(4c\beta\gamma\zeta\mu(\pi_I + \pi_T)(\gamma\zeta(\pi_I + \pi_S + \pi_T) + \pi_N(\mu + \tau)) +$$
$$\left(-\beta c\gamma\zeta(\pi_I + \pi_S + \pi_T) + \mu(\gamma\zeta(\pi_I + \pi_T) + \pi_N(\tau + \phi)) + \mu^2\pi_N\right)^2)$$

$$\tilde{f} = \pi_N\epsilon(\mu + \tau)$$

$$\tilde{g} = 2\beta c$$

$$\tilde{h} = \gamma\zeta(\pi_I + \pi_S + \pi_T) + \pi_N(\mu + \tau)$$

One can also explicitly construct the *fraction* of infected and uninfected needles among the total population of needles at equilibrium by simply dividing either of respective formulae above by $\pi_N/k$. A graph of the fraction of infected needles in the population is given in fig. S6(c).

Let us now consider the effects of small perturbations $\delta k$, around a given value of $k$, on the values of $N_i^*$ and $N_u^*$. We find from a Taylor expansion of $N_i^*(k)$ and $N_u^*(k)$ about $k$ that:

$$N_i^*(k + \delta k) \approx N_i^*(k) - \frac{1}{k}(N_i^*(k) - B(k))\delta k$$
$$N_u^*(k + \delta k) \approx N_u^*(k) - \frac{1}{k}\left(N_u^*(k) + B(k)\right)\delta k$$

where,

$$B(k) = \frac{2\tilde{b}\sqrt{\tilde{c} + \tilde{d}k + \tilde{e}k^2} + \tilde{d} + 2\tilde{e}k}{2\tilde{g}(\tilde{h}k + \tilde{f})\sqrt{\tilde{c} + \tilde{d}k + \tilde{e}k^2}}$$

which can be verified using the fact that $B$ can also be expressed as

$$B(k) = N_i^*(k) + k\frac{dN_i^*}{dk}$$

or by

$$B(k) = -N_u^*(k) + k\frac{dN_u^*}{dk}$$

this follows from the Taylor expansions above—i.e. the term scaling $\delta k$ is $\frac{dN_i^*}{dk}$ in the first equation and $\frac{dN_u^*}{dk}$ in the second equation. Note that $\frac{dN_u^*}{dk}$ follows easily from the expression for $\frac{dN_i^*}{dk}$ since $N_u^* = \frac{\pi_N}{k} - N_i^*$. Evidently, the peak in figure S6(b) occurs because $k$ is such that $N_i^*(k) = B(k)$ ($k \approx 0.5$). This offers some intuition into the meaning of $B(k)$. For a given discard rate $k$, $B(k)$ represents a threshold value for number of infected needles at equilibrium: above $B(k)$, the number of infected needles at equilibrium can be decreased by increasing $k$, whereas when the number of infected needles at equilibrium is below $B(k)$, it can actually increase the number of infected needles at equilibrium to increase $k$. Figure S6(c) is counter-intuitive because it says that *increasing* the discard rate of needles $k$ actually leads to an increase in the equilibrium value of the infected needle fraction. To examine the details of this peculiarity more precisely we look at the equilibrium of the infected needle fraction, which we will denote by $n_i^*(k) = N_i^*/(N_i^* + N_u^*)$.

Using the Taylor expansions from above, it is straightforward to calculate the perturbations about $k$ for the fraction of infected needles. Denoting this fraction by $n_i^*(k)$, we have that

$$n_i^*(k+\delta k) = \frac{N_i^*(k+\delta k)}{N_i^*(k+\delta k)+N_u^*(k+\delta k)} \approx \frac{N_i^*(k)-(N_i^*(k)-B(k))\delta k/k}{(N_i^*(k)+N_u^*(k))\left(1-\frac{\delta k}{k}\right)}$$

and using $N_i^*(k)+N_u^*(k) = \pi_N/k$ along with the approximation $\frac{1}{1-x} \approx 1+x$ with $x = \delta k/k$, we can write

$$\begin{aligned}
n_i^*(k+\delta k) &\approx \frac{k}{\pi_N}\left(N_i^*(k)-(N_i^*(k)-B(k))\left(\frac{\delta k}{k}\right)\right)\left(\frac{1}{1-\frac{\delta k}{k}}\right) \\
&\approx \frac{k}{\pi_N}\left(N_i^*(k)-(N_i^*(k)-B(k))\left(\frac{\delta k}{k}\right)\right)\left(1+\frac{\delta k}{k}\right) \approx n_i^*(k)+\frac{k}{\pi_N}B(k)\left(\frac{\delta k}{k}\right) \\
&= n_i^*(k)+\frac{B(k)}{\pi_N}\delta k
\end{aligned}$$

where in expanding out the product of binomials we used $(\delta k/k)^2 \approx 0$. This shows that the function $B(k)$ is just $\frac{dn_i^*}{dk}$, up to a scaling factor of $\pi_N$. The sign of $B(k)$ determines whether the infected needle fraction (at equilibrium) will increase or decrease with the discard rate $k$. Thus, the number of infected needles in a population will decrease with increasing $k$ if the number of infected needles in the steady state exceeds $B(k)$, which as we have seen is the rate at which the fraction of infected needles increases scaled by $\pi_N$. For the parameters chosen in our model, the value of $B(k)$ is positive for all $k$ values examined (between 0 and 1).

#### *Analytic calculation of* $\mathbf{R_0}$

Below, we provide the $F$ and $V$ matrices used to calculate $R_0$. In this text, we follow the lines of [6], which establishes an algorithm for calculating $R_0$ as the maximum eigenvalue—also called the *spectral radius*—of the matrix $F \cdot V^{-1}$. This matrix is sometimes cited as the *Next Generation Matrix*, however, as Diekmann et. al. (2009) [6] point out, this matrix may be larger than the true next generation matrix. Nonetheless, the maximum eigenvalue of $F \cdot V^{-1}$ will be the same as that of the true next generation matrix. The steps to construct $F$ and $V$ are outlined in the cholera section above. For the case of HCV one finds the following.

$$F = \begin{pmatrix} 0 & 0 & \frac{c\beta k_u}{\mu}\frac{\pi_S}{\pi_N} \\ 0 & 0 & 0 \\ \gamma\zeta & \gamma\zeta & 0 \end{pmatrix}$$

$$V = \begin{pmatrix} \omega+\tau+\mu+\phi & 0 & 0 \\ -\omega & \mu+\tau+\phi & 0 \\ 0 & 0 & \epsilon+k_i \end{pmatrix}$$

From this one can construct $F \cdot V^{-1}$:

$$F \cdot V^{-1} = \begin{pmatrix} 0 & 0 & \frac{c\beta k_u \pi_S}{\mu(\epsilon+k_i)\pi_N} \\ 0 & 0 & 0 \\ \frac{\gamma\zeta}{\mu+\tau+\phi} & \frac{\gamma\zeta}{\mu+\tau+\phi} & 0 \end{pmatrix}$$

From this, one can calculate the maximum eigenvalue of $F \cdot V^{-1}$. One finds,

$$\mathrm{R}_0 = \sqrt{\frac{c\gamma\zeta\beta k_u \pi_S}{\pi_N \mu(\mu+\tau+\phi)(\epsilon+k_i)}}$$

The reproductive ratio is generally interpreted as the average number of secondary infections per capita caused by infected hosts in the time that these hosts have the infection, when the system is near the *disease-free* equilibrium (DFE). In the context of the WAIT modelling scheme, the interpretation of the reproductive ratio may be modified somewhat as the spread of infection is mediated through interactions between living hosts and an environmental intermediate. In this framework, does the $R_0$ value represent the number of new infections within the environment or within the population of living hosts? It will turn out that the $R_0$ value in the WAIT framework represents a kind of average of both of these interpretations, or more precisely, a geometric mean of the two. To put this concretely, we look at the HCV $R_0$ formula from a perspective that illuminates its nature as a geometric mean.

There are two modes of transmission of new infection in the HCV model. One is the infection of clean needles due to injection by infected hosts, and two is the infection of susceptible hosts due to injection by infected needles. These two modes of transmission can each have a reproductive ratio associated with them: the first would be the number of new infected needles caused by infected hosts in the average amount of time that a host is infected (per infected host), and the second would be the number of new infections of susceptible hosts that occur in the average time a needle spends in the infected state, (per infected needle). The first of these two, which we will denote by $X$, can be derived by considering the rate of transmission of infection to clean needles per infected host. In the dynamical equations the total rate is given by $\gamma\zeta(I_E + I_L)\frac{N_u}{N_i+N_u}$, hence the rate *per infected host* is simply $\gamma\zeta\frac{N_u}{N_i+N_u}$. Lastly, at the disease-free equilibrium, the fraction $\frac{N_u}{N_i+N_u}$ is unity since there are no infected needles in the population. This leaves $\gamma\zeta$ as the per capita rate of new infection of needles. Now, the average time that a host is infected (in either of the infected stages, early or late) is simply the reciprocal of the exit rates of the pair of $I_E$ and $I_L$ compartments. That is to say if the average lifetime of an agent within a set, such as the infected hosts, is 5 days, for example, then one would expect 1/5 of the set to leave daily as one expects that on any day 1/5 of the set is comprised of agents who entered the set five days ago—here the assumption is that the set of lifetimes in the set exhibits little variance. That is, the average lifetime within a set is the reciprocal of the exit rate—the exit rate taken as a proportion of the total population, as opposed to a total rate of change in the population size. In the case of the infected population in the HCV model, the exit rates are given by $\mu$,$\tau$, and $\phi$, which are death rate, transfer to treatment rate, and the self-clearance rate, respectively. Thus, the average lifetime of agents within the infected group is given by $1/(\mu+\tau+\phi)$. $X$ is therefore given by $X = \gamma\zeta/(\mu+\tau+\phi)$.

The other reproductive ratio $Y$ can be derived similarly by considering the rate of new host infections due to infected needles: $c\beta S\frac{N_i}{N_i+N_u}$. First, in the disease free equilibrium the value for $S^*$ is given by $\pi_S/\mu$. Second, near the disease-free equilibrium $N_i << N_u$ and hence we can approximate the fraction: $\frac{N_i}{N_i+N_u} = \frac{N_i}{N_u}\frac{1}{\frac{N_i}{N_u}+1} \approx \frac{N_i}{N_u}(1-\frac{N_i}{N_u}) \approx \frac{N_i}{N_u}$, neglecting the $\left(\frac{N_i}{N_u}\right)^2$ term. The result is that near the DFE, the rate term $c\beta S\frac{N_i}{N_i+N_u}$ can be expressed as $c\beta\left(\frac{\pi_N}{\mu}\right)\frac{N_i}{N_u}$. The DFE value for $N_u$—the uninfected needles—is given by $\pi_N/k_u$, which is easily verified by inspection of the $N_u$ dynamical equation. Thus, inserting this into the above rate, and considering the rate *per infected needle* $N_i$, one is left with $c\beta\left(\frac{\pi_N}{\mu}\right)\left(\frac{k_u}{\pi_N}\right)$ as the rate of new host infections caused by infected needles per infected needle. As in the prior argument, the average lifetime a needle spends infected is given simply by the reciprocal of the exit rate for agents in the set. In this case, the exit rate of infected needles is given by $\epsilon$—the virus decay rate—and by $k_i$—the infected needle discard rate. Hence, the average lifetime for an infected needle is given by $1/(\epsilon+k_i)$. Thus, $Y$ is given by $c\beta\left(\frac{\pi_N}{\mu}\right)\left(\frac{k_u}{\pi_N}\right) \times 1/(\epsilon+k_i) = \frac{c\beta\pi_N k_u}{\mu(\epsilon+k_i)\pi_N}$. One will notice that the equation above for $R_0$ can be written as

$$R_0 = \sqrt{\frac{\gamma\zeta}{\mu+\tau+\phi}} \times \sqrt{\frac{c\beta k_u \pi_S}{\mu(\epsilon+k_i)\pi_N}} = \sqrt{XY}$$

In other words, the $R_0$ can be viewed as the geometric mean of the two quantities $X$ and $Y$ discussed above.

#### $\mathbf{R_0}$ *Sensitivity*

We employ a sensitivity analysis for two purposes: (i) to establish that model dynamics do not rely on the particular values of any one parameter and (ii) that the sensitivity of the dynamics is *shared* fairly evenly among the parameters. This is an indication that when the phenomenon of interest are modelled in this manner, one can expect that the dynamics of the infection are also shared fairly evenly among the parameters.

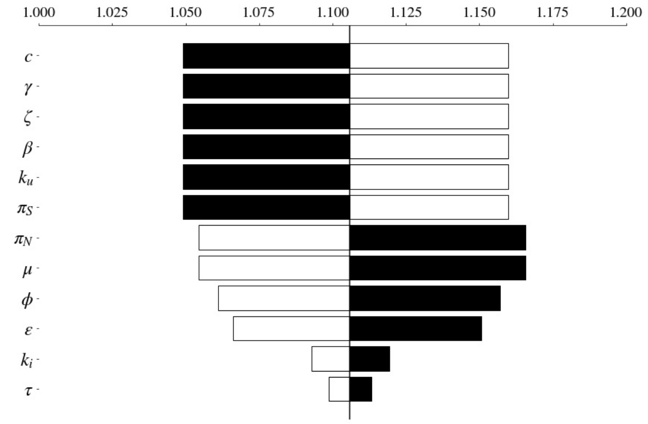


Fig. S7: $\mathbf{R_0}$ **Tornado**: A tornado diagram for parameters in the $R_0$ formula. Black bars indicate the value of $R_0$ when the associated parameter is *decreased* by 10% from the value chosen in the model. White bars indicate the value of $R_0$ when the associated parameter is *increased* by 10%.

Fig. S7 demonstrates how the value of $R_0$ *shares* its dependence evenly among most of the parameters. We calculated the *Partial Rank Correlation Coefficient* (PRCC) with respect to the value of $R_0$, for each of the parameters in the $R_0$ formula (equation 14 in the main text) (Figure 6 in the main text). The calculation followed the lines given in Blower and Dowlatabadi (1994) [7], and we will briefly recapitulate the calculation here.

There are 12 parameters in the $R_0$ formula (equation 14 in the main text). Using Latin Hypercube Sampling (LHS), 100 samples of the 12 parameters were taken. Each sample of 12 parameters was used to calculate a value of $R_0$. Then, the resulting $R_0$ values and parameters were ranked according to their value among the 100 samples. That is, the $R_0$ value and the 12 parameters in each sample were assigned numbers 1–100 depending on how they ranked compared to other samples. This results in 12 vectors, and one additional one for $R_0$, of length 100, whose entries are just some ordering of the whole numbers between 1 and 100—we will call them *rank* vectors. Then, between any two of the 12 rank vectors and one additional rank vector for $R_0$ we can calculate the generic correlation coefficient for the 100 samples. If we arrange the parameters, indexing them 1 through 12, and the $R_0$ value, giving it the index 13, into a list of, let us just generically call them, *variables*, then, we can collect the correlation coefficients $C_{ij}$, between the $ith$ and $jth$ variable, into a symmetric matrix $C$.

$$C_{ij} = \frac{\sum_{k=1}^{100}(r_{ik} - \mu)(r_{jk} - \mu)}{\sqrt{\sum_{k=1}^{100}(r_{ik} - \mu)^2 \sum_{k=1}^{100}(r_{jk} - \mu)^2}}$$

$r_{ik}$ in the equation above is the rank of the $ith$ variable (recall that $R_0$ is included in the variables) in the $kth$ sample, and $\mu = (1 + 100)/2 = 50.5$ is the average rank. Thus, for $i$ and $j$ between 1 and 12, $C_{ij}$ is the correlation coefficient between the $ith$ and $jth$ parameter, and for i between 1 and 12, $C_{i,13}$ $(= C_{13,i})$ is the correlation between the $ith$ parameter and the value of $R_0$. Note, that the diagonal values of $C$ are all one. Next, we construct the matrix $B$, which is simply the matrix inverse of $C$, i.e. $B = C^{-1}$. Lastly, the PRCC value for the $ith$ parameter is constructed from the matrix elements of $B$ according to the following formula,

$$PRCC_i = \frac{-B_{i,13}}{\sqrt{B_{ii}B_{13,13}}}$$

This entire calculation was repeated 50 times, with 50 sets of 100 LHS samples. The values shown in the PRCC calculation (Main text, Figure 6) show the average over these 50 iterations, with the standard deviations as the error bars.

#### References


[1] Richard E Lenski and Robert M May. The evolution of virulence in parasites and pathogens: reconciliation between two competing hypotheses. *Journal of theoretical biology*, 169(3):253–265, 1994.

[2] Paul W Ewald et al. *Evolution of infectious disease*. Oxford University Press on Demand, 1994.

[3] Gandon, Sylvain. The Curse of The Pharoah Hypothesis. *Proceedings of the Royal Society of London B: Biological Sciences*, 265(1405):1545–1552, 1998.

[4] Masashi Kamo and Michael Boots. The curse of the pharaoh in space: free-living infectious stages and the evolution of virulence in spatially explicit populations. *Journal of theoretical biology*, 231(3):435–441, 2004.

[5] Robert I Colautti, Igor A Grigorovich, and Hugh J MacIsaac. Propagule pressure: a null model for biological invasions. *Biological Invasions*, 8(5):1023–1037, 2006.

[6] O. Diekmann, J. A. P. Heesterbeek, and M. G. Roberts. The construction of next-generation matrices for compartmental epidemic models. *Journal of the Royal Society Interface*, 7:873–885, 2010. doi:10.1098/rsif.2009.0386.

[7] S. M. Blower and H. Dowlatabadi. Sensitivity and uncertainty analysis of complex models of disease transmission: An hiv model, as an example. *International Statistical Review*, 62(2):229–243, August, 1994.
